# Phagosome-mediated anti-bacterial immunity is governed by the proton-activated chloride channel in peritoneal macrophages

**DOI:** 10.1101/2025.02.27.640612

**Authors:** Henry Yi Cheng, Jiachen Chu, Nathachit Limjunyawong, Jianan Chen, Yingzhi Ye, Kevin Hong Chen, Nicholas Koylass, Shuying Sun, Xinzhong Dong, Zhaozhu Qiu

## Abstract

The success of phagosome degradation relies on the ability of phagocytes to regulate the maturation of phagosomes. However, its underlying molecular mechanisms remain poorly understood. Here, we identify the proton-activated chloride (PAC) channel as a key negative regulator of phagosome maturation. PAC deletion enhanced phagosomal acidification and protease activities, leading to augmented bacterial killing in large peritoneal macrophages (LPMs) upon *Escherichia coli* infection in mice. Surprisingly, phagosome degradation also stimulated STING-IRF3-interferon responses and inflammasome activation in LPMs, both of which are enhanced upon PAC deletion. The increased inflammasome activation induced the release of cleaved gasdermin D, which localized to the surface of bacteria in the peritoneum and further contributed to their killing. Finally, enhanced bacterial clearance by PAC-deficient LPMs reduced proinflammatory immune cell infiltration and peritoneal inflammation, resulting in improved survival in mice. Our study thus provides new insights into the molecular mechanism of phagosome maturation and the dynamics of host defense response following phagosome-mediated bacterial degradation in peritoneal macrophages.

**Summary:** The PAC channel mediates phagosome maturation during bacterial infection in macrophages. PAC deletion promotes phagosome-mediated STING-interferon signaling and inflammasome-mediated gasdermin D secretion during bacterial infection in peritoneal macrophages.

## Introduction

Phagocytosis represents a fundamental cellular process through which eukaryotic cells internalize large (>0.5 µm) particles, such as microbes, cell corpses and debris, for eventual degradation (Kinchen and Ravichandran, 2008). This process is the first-line host defense in phagocytes, playing a pivotal role in combating various pathogenic infections (Ince et al., 1987). The effectiveness of this defense mechanism hinges on the ability of phagocytes to exert control over the phagosome maturation process, most importantly, through regulating acidity (Beertsen et al., 2008). Upon budding from the plasma membrane, nascent phagosomes undergo a series of maturation steps, acquiring essential components for their acidification. The vesicular proton pump V-ATPase utilizes the energy derived from ATP hydrolysis to transport protons (H^+^) into the phagosomal lumen (Lukacs et al., 1990). Such vectorial transport of H^+^ generates a transmembrane electrical potential difference that would restrain the activity of the pump if left uncompensated. Chloride (Cl^−^) is the major counterion responsible for shunting this luminal positive membrane potential, thus facilitating vesicular acidification (Di et al., 2006). The CLC transporters mediate the accumulation of Cl^−^ within endo-lysosomes, neutralizing the positive charge gradient through their 2Cl^−^ to 1H^+^ exchange activity (Graves et al., 2008; Li et al., 2002). However, their precise contributions to phagosomal acidification and maturation have remained a subject of debate. For example, CLC-3, an early endosomal Cl^−^ exchanger, was linked to phagosomal Cl^−^ accumulation and acidification, but this role was not reported in other studies (Li et al., 2002; Moreland et al., 2006; Wu et al., 2023). Emerging evidence suggests that the phagosomal Cl^−^ concentration itself may also regulate various protease activities. For instance, CLC-7, a lysosomal Cl^−^ exchanger, has been shown to enhance protease activities in phagosomes and promote their fusion with lysosomes, without affecting steady-state phagosomal pH (Wong et al., 2017; Wu et al., 2023). These studies highlight the multifaceted roles of Cl^−^ beyond electrical balancing.

In addition to mediating the killing of microorganisms, phagosomes are also a signaling hub important for innate and adaptive immune responses (Goodridge et al., 2011; Underhill and Goodridge, 2012). For example, innate immune receptors and associated signaling proteins are localized to maturing phagosomes, exhibiting various microorganism-sensing features (Husebye et al., 2010). The antigens of digested pathogens are also processed and interact with the major histocompatibility complex (MHC-I) within the phagolysosome compartments (Nair-Gupta et al., 2014). While most studies focus on phagosomal membrane-bound receptors, emerging evidence suggests that the phagosomal contents, such as peptidoglycan from the gram-positive bacterial cell wall, may also be released after digestion and activate the cytoplasmic pattern recognition receptors (PRRs) (Wolf et al., 2016). However, how the phagosome-mediated degradation and release of bacterial components shape innate immune responses *in vivo* remain underexplored.

Through unbiased RNA interference screens, a novel acid- or proton-activated Cl^−^ (PAC) channel encoded by *PACC1*, also known as *TMEM206*, has been recently identified as a plasma membrane and endosomal Cl^−^ channel (Osei-Owusu et al., 2021; Ullrich et al., 2019; Yang et al., 2019). By releasing Cl^−^ from endosomal lumen under increasing acidic environment, the PAC channel prevents hyperacidification, thus playing an opposite role to the endo-lysosomal CLC exchangers. Notably, PAC is also localized to macropinosomes, facilitating their shrinkage by mediating Cl^−^ efflux (Zeziulia et al., 2022). Despite the importance of Cl^−^, it remains unknown if the PAC channel plays a role in phagosome maturation and how it may contribute to pathogen degradation and subsequently shape the host defense response against infections.

In this study, we identify the PAC channel as an important negative regulator for phagosome maturation in macrophages. Using a peritoneal *Escherichia coli* infection model, we revealed a critical role for PAC in bacterial degradation in large peritoneal macrophages (LPMs). The enhanced bacterial killing in PAC-deficient LPMs activated STING-IRF3-IFNβ axis and inflammasomes through the increased release of bacterial ligands from phagosomes, which further drive the macrophage bactericidal activity. With a novel depletion-adoptive transfer model, we demonstrated crucial roles of PAC and inflammasome-activated gasdermin D in LPM-mediated bacterial clearance. Moreover, we determined how the resolution of peritoneal infection was facilitated by the augmented phagosome bacterial degradation in PAC-deficient LPMs through comprehensive and unbiased analyses, including secretome, RNA-seq, and high-dimensional mass cytometry. Our findings therefore provide new insights into the intricate mechanisms underlying phagosome maturation upon bacterial infection and the dynamics of the innate immune responses triggered by pathogen degradation.

## Results

### The PAC channel localizes to phagosomes and regulates bacterial clearance in macrophages

To identify ion channels and transporters that regulate phagosome maturation, we analyzed a published proteome dataset on phagosomes isolated from mouse macrophages (Guo et al., 2015b) and generated a list of 48 potential candidates (Fig. 1 A and Fig. S1 A). We further cross-referenced with the GeneCards database and narrowed down to 22 proteins that also localize to endo-lysosome compartments thus potentially involved in phagosome maturation. We then mapped the mRNA expression levels of these genes across immune cells with the ImmGen database (Heng et al., 2008). Interestingly, *Pacc1*, encoding the recently identified proton-activated Cl^−^ channel (Ullrich et al., 2019; Yang et al., 2019), was among the highest and most specifically expressed genes across the four types of macrophages (Fig. 1 B and C), including splenic macrophages which are specialized phagocytes relying on phagosomes to degrade their cargo (Flannagan et al., 2009). The proteome and expression analyses suggest a role for the PAC channel in phagosome maturation and macrophage functions. To validate PAC’s phagosomal localization, we performed immunofluorescence analysis on human THP1-differentiated macrophages with a specific monoclonal antibody for human PAC protein. In the resting state, PAC was predominantly localized to early and recycling endosomes, consistent with previous findings in other cell types (Osei-Owusu et al., 2021; Zeziulia et al., 2022) (Fig. S1 B and C). To examine various phagocytosis processes, we applied different substrates, including inorganic silica beads, zymosan particles, and live GFP-expressing *E. coli* (*E. coli*-GFP), and observed the localization of PAC to phagosomes with all three types of cargo (Fig. 1 D and E and Fig. S1 D). PAC was also partially colocalized with LAMP1 in zymosan-containing phagosomes, further confirming its phagolysosome localization (Fig. S1 E). Live cell imaging of Raw264.7 macrophages expressing PAC-mCherry also revealed the association of PAC with the dynamic internalization of *E. coli-GFP* (Fig. 1 F and Fig. S1 F). These data are consistent with the phagosome proteome dataset and a previous report on mouse bone marrow-derived macrophages (BMDMs) treated with dead *E. coli* particles (Zeziulia et al., 2022).

**Figure 1.**
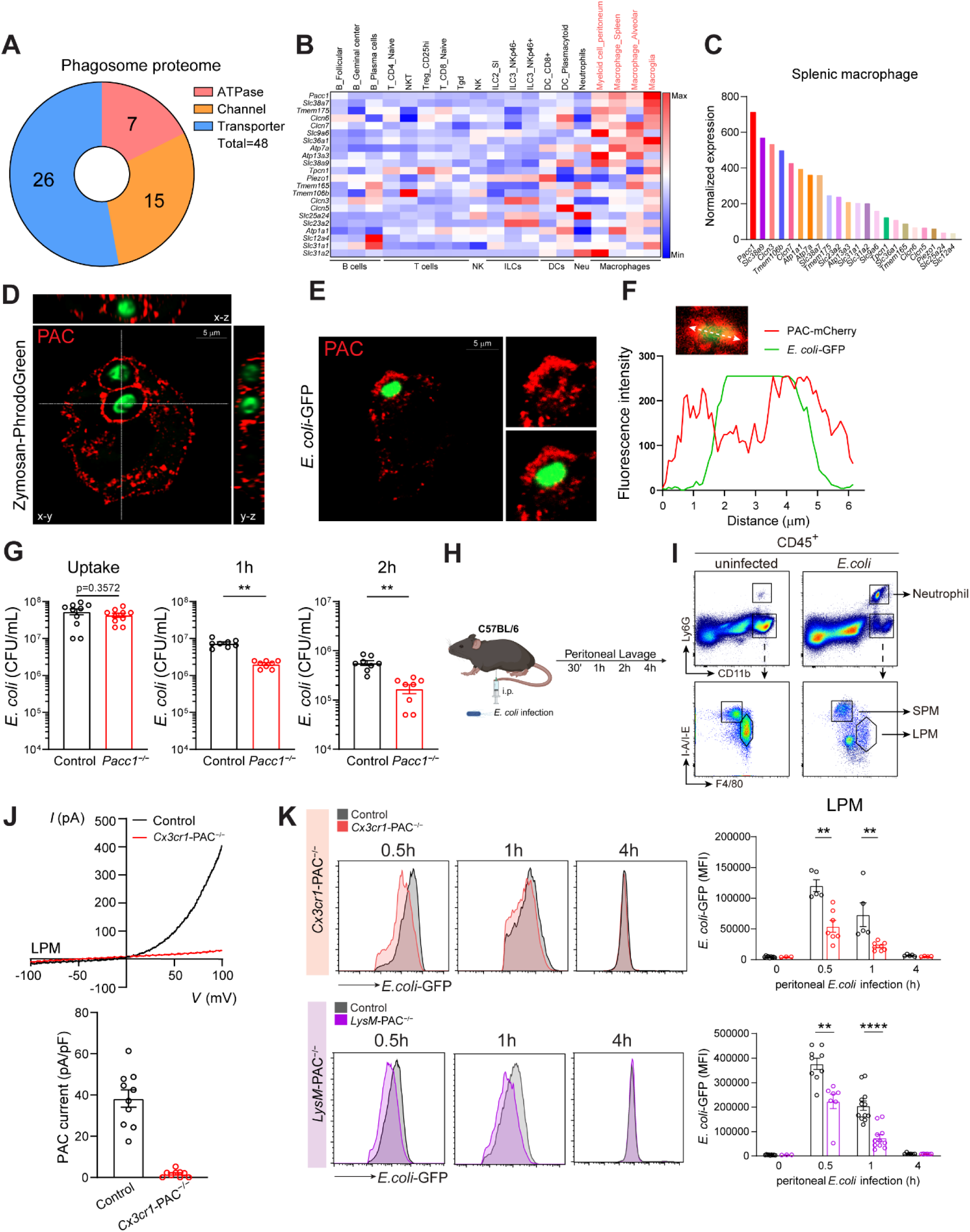
PAC is a phagosome chloride channel mediating macrophage bactericidal activity. **A,** Ion channels and transporters within phagosome proteome (ProteomeXchange: PXD001293). **B,** Gene expression heatmap of different immune cells analyzed in ImmGen. **C,** Gene expression from (**B**) in splenic macrophages. **D-E,** Representative confocal images of THP1 cells with pH-sensing zymosan in z-stacks (**D**) or *E. coli*-GFP in single-plane (**E**) (green) and PAC (red). Data shown as representative of at least 20 cells. Scale bar as indicated. **F**, Fluorescence intensities from confocal live-cell imaging of PAC-mCherry and infected *E. coli*-GFP at 9 min in Fig. S1F. Data shown as representative of at least 20 cells. **G**, BMDM intracellular bacterial culture. Data represented from n = 8 independent experiments. **H**, Schematic illustration of peritoneal infection model. **I**, Gating strategy for peritoneal neutrophils, LPM, and SPM. **J**, Representative I-V relationship (top) and current density (bottom) of whole-cell PAC current induced by pH 4.6. control, n = 10 cells; *Cx3cr1*-PAC^−/−^, n = 9 cells. **K**, FACS analysis histogram (left) and MFI quantification (right) of *in vivo E. coli*-GFP clearance assay. For histograms, *E. coli*-GFP fluorescence in LPM was plotted at indicated times. For quantifications, GFP MFI in LPM was quantified at indicated times. At least 500,000 cells were collected per condition. Data represented from n > 4 mice. All control groups represent *Pacc1^F/F^* genotype. Data are reported as mean ± SEM between independent experiments. Unpaired t-test for **G**. Two-way ANOVA with Sidak’s test for **K**. **p < 0.01, ****p < 0.0001.

To further examine the function of PAC in phagosome maturation and macrophage functions, we cultured BMDMs from PAC whole-body knockout (*Pacc1*^−/−^; KO) mice and validated the loss of PAC-specific acid-induced Cl^−^ currents (*I_Cl, H_*) by patch clamp recordings (Fig. S1 G). PAC deletion did not alter BMDM differentiation from bone-marrow hematopoietic stem cells treated with either L929 cell conditioned media or M-CSF supplement (Fig. S1 H). Given the critical role of macrophages in clearing bacterial infection, we next evaluate the bactericidal activity of *Pacc1*^−/−^ BMDMs by infecting them with *E. coli*. Interestingly, while *Pacc1*^−/−^ BMDMs internalized a similar number of *E. coli* as control cells, the surviving bacteria within the KO cells were less than WT cells after 1-2 hours of digestion, indicating an enhanced bactericidal activity in macrophages without PAC channel (Fig. 1 G).

To further examine the role of PAC in bacterial clearance *in vivo*, we adopted a peritoneal *E. coli* infection model (Fig. 1 H). In this model, immune cells can be obtained from peritoneal lavage immediately after infection and subjected to fluorescence-activated cell sorting (FACS) analyses and functional studies. Large peritoneal macrophages (LPMs) and small peritoneal macrophages (SPMs) are two resident macrophages in the peritoneum, characterized as the CD11b^+^, F4/80^high^, and I-A/I-E^low^ population, and CD11b^+^, F4/80^low^, and I-A/I-E^high^ population, respectively (Fig. 1 I). LPMs in particular are the major phagocytes of the first-line defense against peritoneal infection (Vega-Perez et al., 2021). To examine the role of PAC in bacterial killing, we crossed *Pacc1^F/F^*mice with *Cx3cr1* promoter-driven Cre mice to selectively delete the gene in all resident macrophages and monocytes (*Cx3cr1*-PAC^−/−^). The loss of PAC channel activity in LPMs isolated from *Cx3cr1*-PAC^−/−^ mice was validated by patch clamp recordings (Fig. 1 J). The mutant mice had similar numbers of LPMs, SPMs, and monocytes in the peritoneum compared to their *Pacc1^F/F^* control littermates, suggesting that the PAC channel is not required for their general development (Fig. S2 A and B). Neutrophils and various lymphocyte (CD4 T cells, CD8 T cells, B cells, and NK cells) populations in the peritoneum were also normal (Fig. S2 B-E). For real-time *in vivo* tracking of bacterial clearance, we peritoneally infected mice with *E. coli*-GFP and analyzed GFP intensity in cells isolated from the peritoneal lavage. Since PAC deletion does not affect initial phagocytosis or bacterial uptake in BMDMs (Fig. 1 G), we hypothesized that GFP fluorescence within various immune cells reflects their bactericidal activity. Consistent with the enhanced bacterial clearance *in vitro*, GFP fluorescence declined more rapidly in LPMs from *Cx3cr1*-PAC^−/−^ mice compared to controls (Fig. 1 K). This was specific to LPMs, as neutrophil clearance was similar between groups. No *E. coli*-GFP signals above backgrounds were detected in SPMs, consistent with their non-phagocyte nature (Fig. S2 G). Although acidic pH in phagosomes may also contribute to GFP quenching, reduced GFP fluorescence likely reflects phagosomal bacteriolysis activity as it is required to expose intracellular GFP to acidic phagosomes. Altogether, these results suggest that *Cx3cr1*-PAC^−/−^ LPMs exhibit enhanced bacteriolysis activities.

*Cx3cr1*-Cre line was generated through a knock-in strategy, leading to the deletion of one allele of endogenous *Cx3cr1*, a chemokine receptor (Yona et al., 2013). Although the immune system development was grossly normal in *Cx3cr1*-PAC^−/−^ mice, we further validated our findings by crossing *Pacc1^F/F^* mice with *LysM*-Cre (Clausen et al., 1999), another popular Cre line specific for myeloid cell lineages to generate *LysM*-PAC^−/−^ mice. Consistently, *LysM*-PAC^−/−^ mice also had normal numbers of peritoneal myeloid populations (Fig. S2 A and C) and lymphoid populations (Fig. S2 D and F). We next performed peritoneal infection of *E. coli*-GFP and observed comparable decreases in GFP intensity in LPMs from *LysM*-PAC^−/−^ mice, similar to those from *Cx3cr1*-PAC^−/−^ mice (Fig. 1 K and Fig. S2 G). Taken together, these results from both *in vitro* and *in vivo* experiments with two independent Cre lines demonstrate that PAC-deficient macrophages exhibit an enhanced bacterial clearance capability.

### PAC downregulation is required for LPS-induced phagosome maturation

To further explore the role of PAC in bacterial infections, we analyzed *PACC1* expression in published transcriptome databases of different infectious diseases (Bertrams et al., 2020; Scicluna et al., 2020). Interestingly, *PACC1* was significantly downregulated in the peripheral blood mononuclear cells (PBMCs) from patients with sepsis or pneumonia (Fig. 2 A). We hypothesized that PAC downregulation occurs in mononuclear cells upon pathogen recognition. To test this, we treated BMDMs with lipopolysaccharides (LPS), a key bacterial component that induces immune activation (Poltorak et al., 1998). Strikingly, LPS treatment markedly reduced *Pacc1* mRNA levels as well as PAC channel activity (Fig. 2 B and C). Pretreatment of TAK-242, a Toll-like receptor 4 (TLR4) inhibitor, before LPS stimulation completely abolished the LPS-induced *Pacc1* downregulation (Fig. 2 D), indicating that the regulation of *Pacc1* expression is dependent on the LPS-TLR4 signaling. This regulation appears to be specific to the PAC channel as the activity of another chloride channel, volume-regulated anion channel (VRAC), was unaltered in LPS-treated BMDMs (Fig. S3 A and B). The same LPS-induced PAC downregulation was also observed in human THP1 cells at both mRNA and protein levels (Fig. S3 C and D).

**Figure 2.**
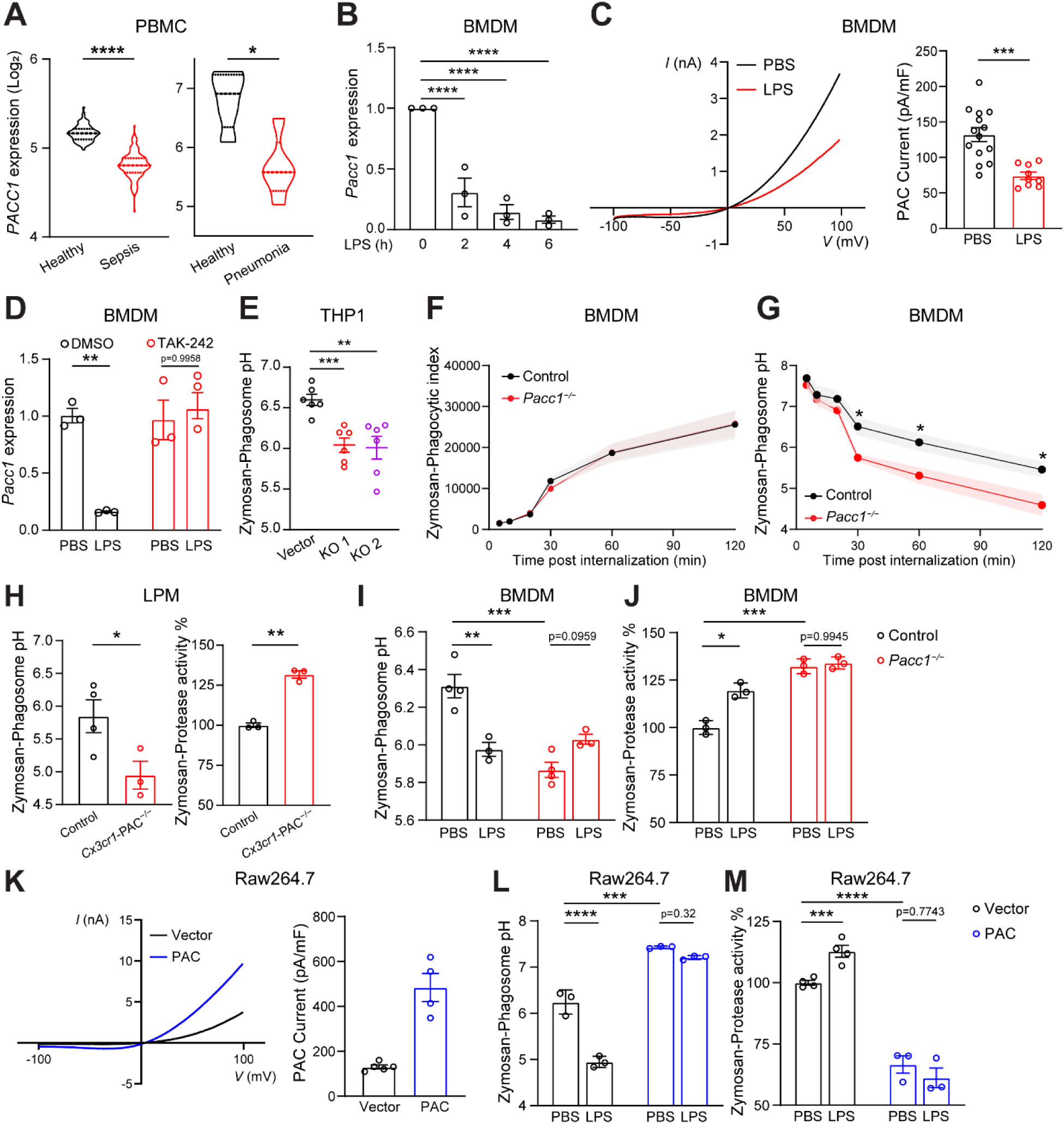
PAC downregulation is required for LPS-induced phagosome maturation. **A**, *PACC1* expression analyzed in GSE134364 (left) or GSE94916 (right). **B**, *Pacc1* mRNA expression in BMDMs after LPS (100 ng/mL) treatments. Data represented from n = 3 mice. **C**, Representative I-V relationship (left) and current density (right) of whole-cell PAC current in BMDMs post 4 hours LPS (1 μg/mL) treatment. PBS: n = 14 cells from 3 mice. LPS: n = 9 cells from 3 mice. **D,** *Pacc1* mRNA expression in BMDMs pretreated with TAK-242 (10 nM) for 30 min followed by 4 hours of LPS (1 μg/mL) treatment. Data represented from n = 3 mice. **E**, FACS measurement of phagosome pH in THP1 cells. Cells were measured 30 min post zymosan internalization. Data represented from n = 6 independent experiments. **F**, FACS measurement of phagocytosis activity in BMDMs. Data represented from n > 4 mice at each timepoint. **G,** FACS measurement of phagosome pH in BMDMs. For treatments longer than 30 min, zymosan was washed out after 30 min of internalization. Data represented from n > 4 mice at each timepoint. **H**, FACS measurement of phagosome pH (left) or protease activity (right) in LPMs 30 min post zymosan peritoneal injection. Data represented from n = 3 mice. **I-J**, FACS measurement of phagosome pH (**H**) or protease activity (**I**) in BMDMs. Cells were measured 30 min post zymosan internalization after 4 hours LPS (20 ng/mL) treatment. Data represented from n > 3 mice. **K**, Representative I-V relationship (left) and current density (right) of whole-cell PAC current in Raw264.7 cells. Vector, n = 5 cells; PAC, n = 4 cells. **L-M**, FACS measurement of phagosome pH (**L**) or protease activity (**K**) in Raw264.7 cells. Cells were measured 30 min post zymosan internalization after 4 hours LPS (20 ng/mL) treatment. Data represented from n > 3 independent experiments. All control groups represent *Pacc1^F/F^* genotype. Data are reported as mean ± SEM between independent experiments. Unpaired t-test for **A**, **C, H**. One-way ANOVA with Fisher’s LSD test for **B, E**. Two-way ANOVA with Sidak’s test for **D**, **G, I, J, L, M**. *p < 0.05, **p < 0.01, ***p < 0.001, ****p < 0.0001.

Given its role in vesicular acidification (Osei-Owusu et al., 2021; Zeziulia et al., 2022), we tested whether the PAC channel regulates phagosomal acidification by generating PAC KO THP1 cells using CRISPR-Cas9 approach (Fig. S3 E). We next performed ratiometric measurement of phagosomal pH using zymosan particles conjugated with pH-sensitive PhrodoGreen and pH-insensitive Alexa Fluor-633 (AF633) fluorescence dyes. Interestingly, loss of PAC resulted in ∼0.5 unit lower pH in phagosomes at steady-state compared to WT THP1 cells (Fig. 2 E). We further validated this finding in BMDMs and LPMs. Consistent with *E. coli* uptake results (Fig. 1 G), *Pacc1*^−/−^ BMDMs also had a normal uptake of zymosan particles (Fig. 2 F). However, phagosomes were more acidic during the maturation process in the KO BMDMs compared to WT control cells (Fig. 2 G). Additionally, we observed a similar decrease in phagosomal pH in LPMs isolated from *Cx3cr1*-PAC^−/−^ mice after peritoneally injecting pH-sensing zymosan particles (Fig. 2 H, left). Given that the activities of phagosomal proteases are dependent on acidic pH (Kinchen and Ravichandran, 2008), we measured them by a similar ratiometric method using zymosan particles double-conjugated with bovine serum albumin (BSA) self-quenched DQ green (BSA-DQ) fluorescence dye and AF633. The protease activities were indicated by the ratio of the green fluorescence detected after BSA is degraded by phagosomal proteases to internal control AF633. Consistent with enhanced phagosomal acidification, we also observed a modest but significant increase in protease activities in LPMs isolated from *Cx3cr1*-PAC^−/−^ mice after peritoneally injecting BSA-DQ zymosans (Fig. 2 H, right). Together, these results demonstrate that the PAC channel plays a negative role in phagosome acidification and maturation.

LPS-primed macrophages exhibit enhanced phagosome degradation, suggesting that LPS promotes phagosome acidification and maturation (Wu et al., 2023). Indeed, BMDMs primed with LPS had more acidic phagosomes with increased overall protease activities (Fig. 2 I and J, black). Without LPS treatment, *Pacc1*^−/−^ BMDMs already exhibited a similarly more acidic phagosomal pH and higher protease activities at the baseline (Fig. 2 I and J, red), indicating that loss of PAC mimics the effect of LPS on phagosome maturation. Importantly, treating *Pacc1*^−/−^ BMDMs with LPS did not generate further responses, suggesting that the PAC channel may mediate the effect of LPS priming on phagosome maturation. To test this hypothesis, we overexpressed the PAC protein in Raw264.7 macrophages (Fig. 2 K). Consistent with a negative role in phagosome maturation, PAC overexpression alkalized phagosomal pH and inhibited overall protease activities (Fig. 2 L and M). Similar to BMDMs, LPS priming promoted phagosome activation in vector-transfected Raw264.7 macrophages (Fig. 2 L and M, black). However, these effects were abolished in PAC-overexpressing cells (Fig. 2 L and M, blue), further suggesting that PAC downregulation is required for LPS-mediated phagosome activation. In addition to phagosome maturation, LPS also activates macrophage TLR4 and its downstream IRF3 and NFκB signaling pathways (Poltorak et al., 1998). Interestingly, PAC deletion did not alter LPS-induced expression of IRF3-regulated *IFNβ* and NFκB-mediated transcripts (*IL-6*, *TNFα*, and *IL-1β*) in THP-1 and BMDMs (Fig. S3 F). These results indicate that PAC does not regulate LPS-induced TLR4 activation.

### Enhanced phagosome bacterial degradation promotes interferon responses in PAC-deficient LPMs

To further investigate potential downstream responses following enhanced bacterial degradation, we performed RNA-sequencing (RNA-seq) analysis on LPMs isolated from *Cx3cr1*-PAC^−/−^ and control mice after peritoneal *E. coli* infection. Interestingly, the gene enrichment analysis (GSEA) showed that the signatures of “killing of cells of another organism” and “MHC-I protein binding” were upregulated in PAC-deficient LPMs compared to control cells (Fig. 3 A). Given the antigen processing of MHC-I complexes occurs in phagosomes, these results are consistent with enhanced phagosome function. The gene ontology (GO) analysis further revealed upregulation of interferon responses in PAC-deficient LPMs (Fig. 3 B and Fig. S4 A and B), with increased expression of many IFN-responsive genes, including several *Ifi*, *Gbp*, and *Oas* family genes (Fig. 3 C). To validate the enhanced IFN responses, we infected LPMs with *E. coli in vitro* and performed a comprehensive secretome analysis (Fig. S4 C and D). In accordance with the RNA-seq data, *Pacc1*^−/−^ LPMs showed increased release of IFNβ and some other cytokines compared to control LPMs (Fig. 3 D and Fig. S4 E). Specifically, the KO cells released more chemokines (CXCL1, CCL3, and CCL4) and alarmins (IL-1α, IL-23, and IL-33) (Fig. S4 F and G), which may contribute to LPM activation and host defense against bacterial infection (Chan et al., 2012; Tilstam et al., 2021). Interestingly, we also observed increased releases of IL-1β and IL-18 in *Pacc1*^−/−^ LPMs, indicating enhanced inflammasome activation (Fig. S4 H).

**Figure 3.**
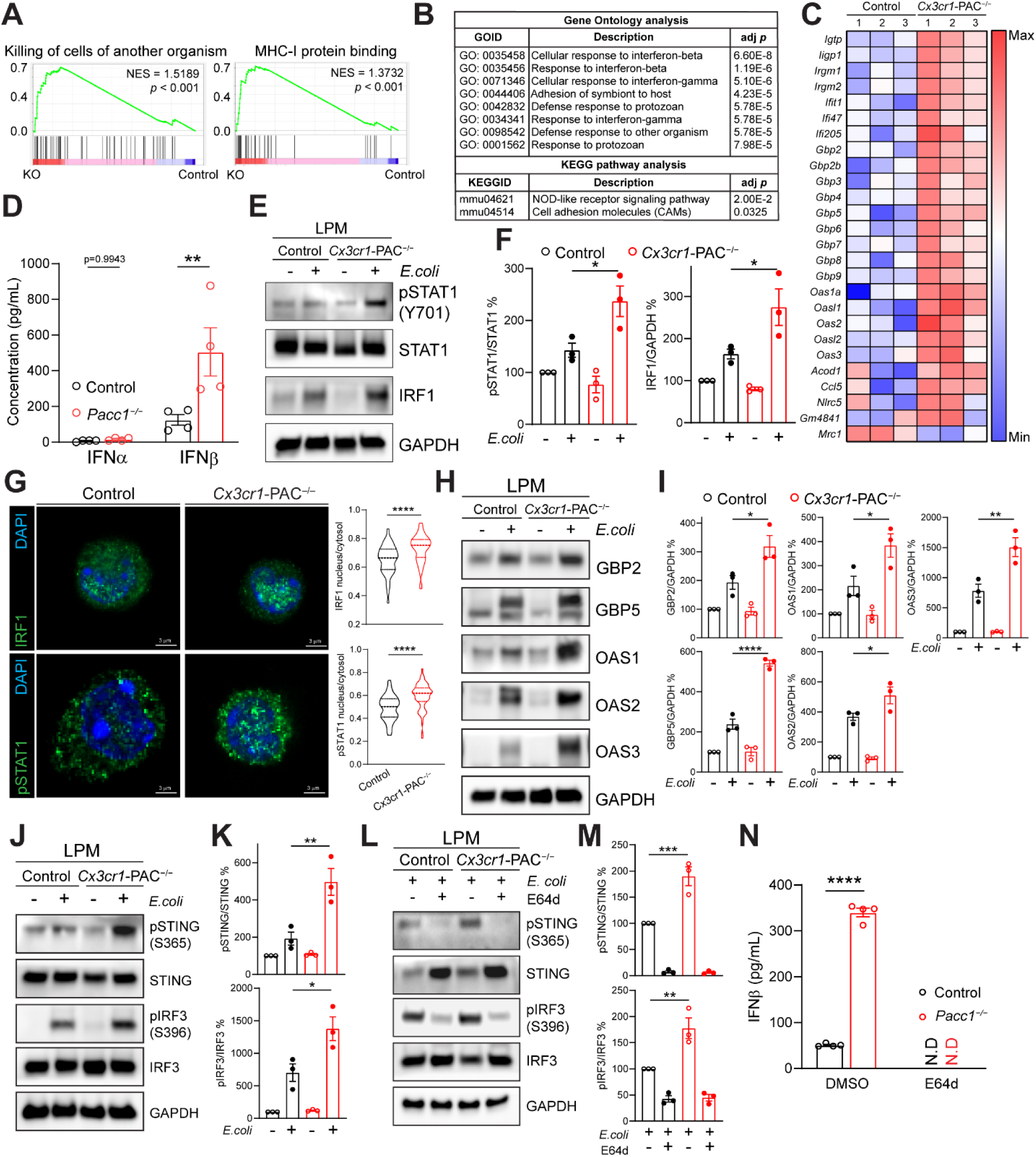
Enhanced phagosome bacterial degradation promotes interferon responses in PAC-deficient LPMs. **A,** GSEA signatures of FACS-sorted LPM RNA-seq analysis performed 30 min post infection. NES, normalized enrichment score. **B**, GO terms and KEGG pathways of FACS-sorted LPM RNA-seq analysis performed 30 min post infection. **C**, Heatmap depicting Z-score of highlighted gene expressions in RNA-seq analysis. **D**, ELISA of IFNα and IFNβ from MACS-isolated LPMs after *E. coli* infection *in vitro*. Data represented from n = 4 mice. **E-F**, Immunoblotting (**E**) and quantification (**F**) of MACS-isolated LPMs. LPMs from 2 mice 60 min post peritoneal infection were pooled and used in 1 experiment. Data represented from n = 3 independent experiments. **G**, Representative confocal images (left) in single-plane and quantification (right) of MACS-isolated LPMs 60 min post peritoneal infection. Control LPM for IRF1, n = 45 cells; *Cx3cr1*-PAC^−/−^ LPM for IRF1, n = 53 cells; control LPM for pSTAT1, n = 90 cells; *Cx3cr1*-PAC^−/−^ LPM for pSTAT1, n = 74 cells. **H-I**, Immunoblotting (**H**) and quantification (**I**) of MACS-isolated LPMs. LPMs from 2 mice 60 min post peritoneal infection were pooled and used in 1 experiment. Data represented from n = 3 independent experiments. **J-K**, Immunoblotting (**J**) and quantification (**K**) of MACS-isolated LPMs. LPMs from 2 mice 15 min post peritoneal infection were pooled and used in 1 experiment. Data represented from n = 3 independent experiments. **L-M,** Immunoblotting (**L**) and quantification (**M**) of MACS-isolated LPMs. LPMs from 2 mice pretreated with E64d (0.15 mg/kg) following 15 min of peritoneal infection were pooled and used in 1 experiment. Data represented from n = 3 independent experiments. **N**, ELISA of IFNβ from MACS-isolated LPMs pretreated with E64d (5 μM) following *E. coli* infection *in vitro*. Data represented from n = 4 mice. N.D, not detected. All control groups represent *Pacc1^F/F^* genotype. Data are reported as mean ± SEM between independent experiments. Unpaired t-test for **D**, **G**. Two-way ANOVA with Sidak’s test for **F, I**, **K, M, N**. *p < 0.05, **p < 0.01, ****p < 0.0001.

With increased IFNβ production in LPMs from *Cx3cr1*-PAC^−/−^ mice after peritoneal *E. coli* infection, we further examined its downstream signaling pathways. The IFNβ-induced phosphorylation of STAT1 and downstream IRF1 expression drive the transcription of IFN responsive genes during bacterial infections (Man et al., 2016). Consistent with RNA-seq and secretome results, immunoblotting revealed increases in STAT1 phosphorylation and IRF1 expression in PAC-deficient LPMs compared to control cells (Fig. 3 E and F). Immunostaining further showed elevations of their translocation into nucleus (Fig. 3 G). The increased expression of downstream GBP and OAS family proteins in *Cx3cr1*-PAC^−/−^ LPMs was also validated by immunoblotting and FACS analyses (Fig. 3 H-I and Fig. S4 I). These data suggest that the increased activation of IFNβ-STAT1-IRF1 axis in PAC-deficient LPMs drives the expression of crucial macrophage anti-microbial genes, such as *Gbp* and *Oas*. (Fisch et al., 2023; Hornung et al., 2014; Leisching et al., 2019; Man et al., 2016).

### Enhanced phagosome bacterial degradation induces IFNβ expression through STING-IRF3 activation in PAC-deficient LPMs

We next investigated the potential mechanism underlying the enhanced IFNβ production in PAC-deficient LPMs upon peritoneal *E. coli* infection. During bacterial infection, the production of IFNβ is regulated by cell surface TLR4 and cytoplasmic pattern recognition receptors (PRRs) (Honda et al., 2006; Manzanillo et al., 2012). Given that no significant change in TLR4-induced genes was observed in *Pacc1*^−/−^ BMDMs upon LPS treatment (Fig. S3 F), we reasoned that the increased IFNβ production in PAC-deficient LPMs may be driven by augmented activation of cytoplasmic PRRs. Interestingly, the STING (stimulator of interferon genes) receptor has been reported to drive macrophage IFN production in various bacterial infections (Andrade et al., 2016; Manzanillo et al., 2012; Moretti et al., 2017; Ruangkiattikul et al., 2017). Therefore, we hypothesized that the enhanced phagosome bacterial degradation in PAC-deficient LPMs drives STING hyperactivation, thereby promoting downstream IFNβ production. To test this hypothesis, we performed immunoblotting and evaluated the phosphorylation of STING and the downstream transcription factor IRF3. Indeed, we observed the activation of STING-IRF3 signaling in control LPMs upon peritoneal *E. coli* infection. More importantly, STING-IRF3 activation was further enhanced in *Cx3cr1*-PAC^−/−^ LPMs (Fig. 3 J and K). We next assayed NFκB p65 activation which is also downstream of STING as well as other cytoplasmic PRRs (Abdullah and Knolle, 2014). Consistently, the phosphorylation and translocation of p65 were also augmented in *Cx3cr1*-PAC^−/−^ LPMs (Fig. S4 J and K). To further examine the role of phagosome degradation in interferon responses, we treated mice with E64d, a phagosome pan-cathepsin inhibitor, which potently blocked the bactericidal activity of LPMs in the peritoneal *E. coli* infection model (Fig. S4 L). E64d treatment also abolished the activation of STING-IRF3 pathway in both control and *Cx3cr1*-PAC^−/−^ LPMs (Fig. 3 L and M), thereby eliminating IFNβ production (Fig. 3 N). Notably, E64d treatment increased total STING protein expression (Fig. 3 L and M), likely due to its effect on STING lysosomal turnover properties (Kuchitsu et al., 2023). Even though STING was stabilized upon E64d treatment, no phosphorylated STING was detected, further demonstrating the dependency of STING activation on phagosome bacterial degradation. Together, our findings provide an underappreciated mechanism downstream of phagosome degradation, in which STING-IRF3 axis is activated by phagosome bacterial degradation alongside the IFNβ-STAT1-IRF1 activation to induce host defense genes expression in LPMs. The enhanced phagosome degradation in PAC-deficient macrophages further promotes the activation loop and creates a hyperactivated status in response to *E. coli* infection.

### Augmented phagosome bacterial degradation induces inflammasome activation and pyroptosis in PAC-deficient LPMs

LPMs have been shown to undergo pyroptosis, but not apoptosis or necroptosis during peritoneal *E. coli* infection (Vega-Perez et al., 2021). Consistently, we observed releases of IL-1β, IL-18, and lactate dehydrogenase (LDH) from control LPMs following *in vitro E. coli* infection, with significantly higher releases in *Pacc1*^−/−^ LPMs compared to controls, suggesting enhanced inflammasome activation and pyroptosis in the absence of PAC (Fig. 4 A and Fig. S4 H). The treatment of necrosulfonamide (NSA), a gasdermin D inhibitor, abolished IL-1β release in both groups of cells, excluding the possibility of inflammasome-independent IL-1β release (Fig. S5 A). We next examined the LPM inflammasome status *in vivo* by isolating and characterizing them after peritoneal *E. coli* infection. Consistently, ∼40% LPMs isolated from control mice were Annexin V and 7-AAD double positive, indicating ongoing cell death. More LPMs underwent cell death in *Cx3cr1*-PAC^−/−^ mice (Fig. 4 B). Accordingly, more gasdermin D pores were observed on their cell surface by non-permeabilized immunostaining (Fig. 4 C and D) and FACS analysis (Fig. S5 B), indicating increased pyroptotic cell death. Inflammasome activation entails caspase-1 (CASP1) cleavage into active CASP1 p20, further leading to the cleavage of gasdermin D into pore-forming N-terminus gasdermin D (gasdermin D-NT) (Barnett et al., 2023). Consistent with increased gasdermin D pore formation, immunoblotting analysis revealed that the cleavage of CASP1 and its downstream target gasdermin D were elevated in LPMs from *Cx3cr1*-PAC^−/−^ mice after peritoneal *E. coli* infection (Fig. 4 E and F). The increased CASP1 activation was also validated by FACS analysis using fluorescein-conjugated CASP1 inhibitor FLICA (FLICA-FAM), which specifically binds to the active CASP1 cleavage site (Fig. S5 C).

**Figure 4.**
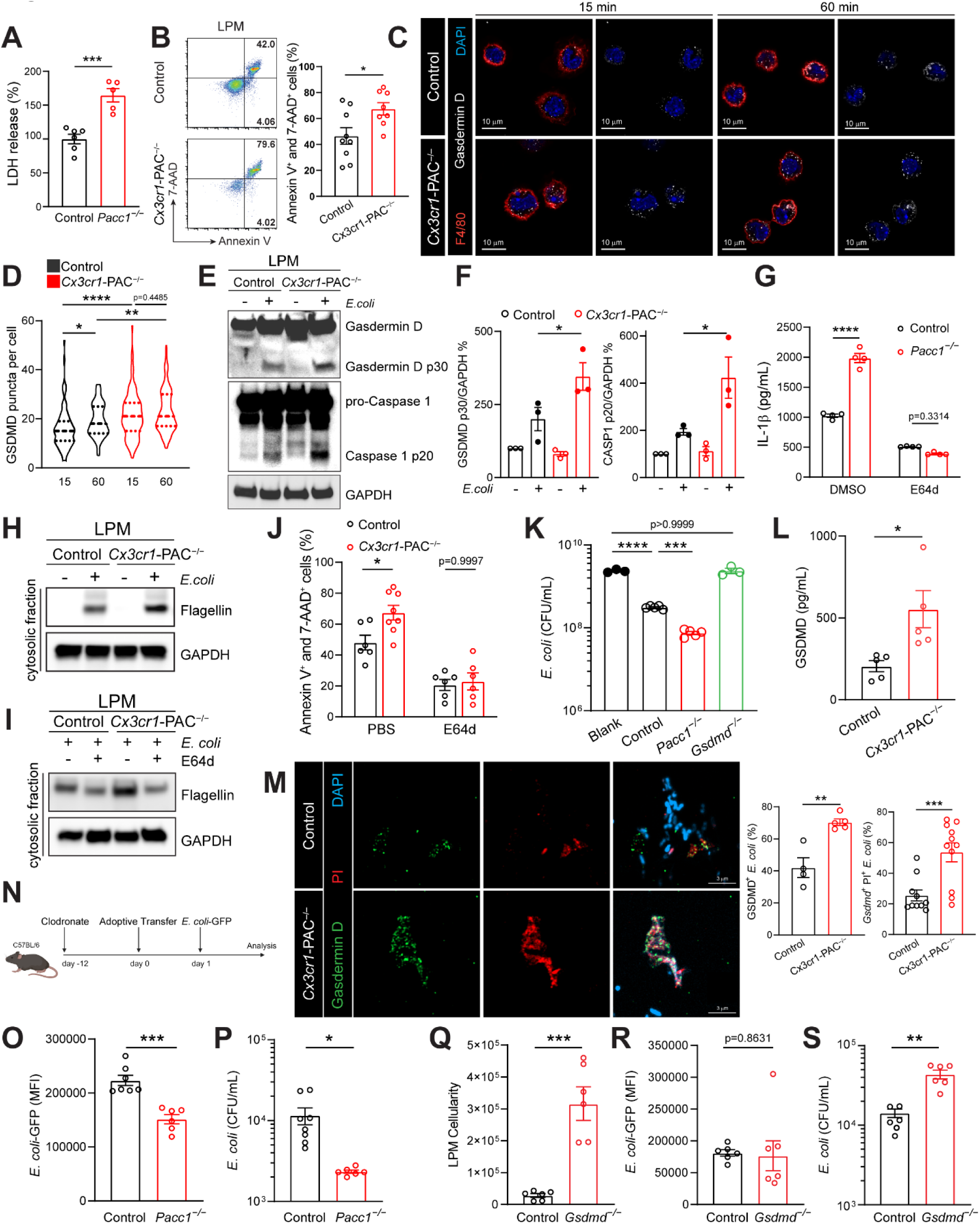
Augmented phagosome bacterial degradation facilitates inflammasome activation and pyroptosis in PAC-deficient LPMs. **A,** Colorimetric LDH assay from MACS-isolated LPMs after *E. coli* infection *in vitro*. Data represented from n = 5 mice. **B,** FACS analysis (left) and quantification (right) of annexin-V^+^ and 7-AAD^+^ LPMs within peritoneal fluids after 30 min of infection. Data represented from n > 8 mice. **C-D**, Representative confocal images (**C**) and quantification (**D**) of MACS-isolated LPMs post peritoneal infection. 15 min control, n = 98 cells; 15 min *Cx3cr1*-PAC^−/−^, n = 109; 60 min control, n = 99 cells, 60 min *Cx3cr1*-PAC^−/−^, n = 108. **E-F**, Immunoblotting (**E**) and quantification (**F**) of MACS-isolated LPMs. LPMs from 4 mice 30 min post peritoneal infection were pooled and used in 1 experiment. Data represented from n = 3 independent experiments. **G**, ELISA of IL-1β from MACS-isolated LPMs pretreated with E64d (5 μM) following *E. coli* infection *in vitro*. Data represented from n = 4 mice. **H-I**, Immunoblotting of MACS-isolated LPM cytosolic extraction. LPMs from 2 mice without treatment (**H**) or pretreated with E64d (0.15 mg/kg) 15 min before peritoneal infection (**I**) were pooled and used in 1 experiment. Data represented from n = 3 independent experiments. **J**, FACS quantification of annexin-V^+^ and 7-AAD^+^ LPMs within peritoneal fluids pretreated with E64d (0.15 mg/kg) before peritoneal infection. Data represented from n > 6 mice. **K**, *E. coli* cultured with antibiotic-free supernatants, collected from *E. coli*-infected control, *Pacc1^−/−^*, or *Gsdmd^−/−^* LPMs, for 6 h at 37C. Blank, antibiotic-free RPMI medium. **L**, ELISA of gasdermin D from peritoneal fluids after 60 min of infection. Data represented from n = 5 mice. **M,** Representative confocal images (left) and quantification (right) of isolated *E. coli* within peritoneal fluids after 60 min of infection. Data represented from n > 4 mice. **N**, Schematic illustration of depletion-adoptive transfer model. **O-P**, FACS quantification of GFP intensity in CellTrace^+^ LPM (**O**) and peritoneal bacterial culture (**P**) within LPM-depleted mice. Data represented from n > 6 mice. **Q-S**, FACS quantification of CellTrace^+^ LPM cellularity (**Q**), GFP intensity (**R**) and peritoneal bacterial culture (**S**) within LPM-depleted mice. Data represented from n = 6 mice. Control represents *Pacc1^F/F^* genotype in **A-P** or C57BL/6 wildtype in **Q-S**. Data are reported as mean ± SEM between independent experiments. Unpaired t-test for **A, B, L, M, O, P, Q, R, S**. One-way ANOVA with Fisher’s LSD test for **K**. Two-way ANOVA with Sidak’s test for **D**, **F, G, J**. *p < 0.05, **p < 0.01, ***p < 0.001, ****p < 0.0001.

To investigate the mechanism underlying the increased inflammasome activation in PAC-deficient LPMs, we first evaluated the IL-1β secretion with two canonical inflammasome activators, ATP and nigericin, after LPS priming. Surprisingly, no difference between control and *Pacc1*^−/−^ LPMs was observed, suggesting that PAC regulates inflammasome activation specifically in the context of bacterial infection, likely involving phagosome degradation (Fig. S5 D, left). To test this hypothesis, we treated LPMs with E64d, which, as expected, had no effect on the canonical ATP-induced IL-1β secretion (Fig. S5 D, right). However, E64d treatment reduced *E. coli*-induced IL-1β release in control LPMs. More importantly, E64d entirely abolished the potentiation of IL-1β release observed in *Pacc1*^−/−^ LPMs (Fig. 4 G). These data suggest that the augmented phagosome bacterial degradation in PAC-deficient LPMs leads to the enhanced inflammasome activation.

Inflammasomes are formed by different sensors that are activated by various cytoplasmic bacterial ligands, such as LPS, flagellin, and bacterial DNA (Barnett et al., 2023). Given that we did not observe *E. coli* escaping from phagosomes, the phagosome degradation-mediated inflammasome activation indicates a potential release of these bacterial ligands from phagosomes to the cytoplasm. Among various ligands, flagellin can be readily detected by antibody-based assays. Therefore, we used it as a readout of released bacterial components. We first performed immunofluorescence analysis on flagellin in LPMs isolated after peritoneal *E. coli* infection. As expected, large flagellin signals were co-localized with LAMP1, indicating bacteria within phagolysosomes. Interestingly, we also observed many small flagellin-positive puncta in the cytoplasm of both control and *Cx3cr1*-PAC^−/−^ LPMs, which did not colocalize with LAMP1^+^ phagolysosomes (Fig. S5 E). Consistently, flagellin proteins were also detected (Fig. 4 H and S5 F). In accordance with enhanced phagosome degradation, the level of cytosolic flagellin was higher in *Cx3cr1*-PAC^−/−^ LPMs compared to control cells (Fig. 4 H and S5 G). More importantly, peritoneal injection of E64d before *E. coli* infection blocked flagellin release in the cytoplasm and eliminated the elevation of cell death in *Cx3cr1*-PAC^−/−^ LPMs (Fig. 4 I-J and S5 H). These results suggest the release of bacterial components after phagosomal degradation. The cytosolic presence of these bacterial components could activate different inflammasome sensors, leading to CASP1 and gasdermin D activation. We examined whether NLRP3, a major bacterial toxin and cellular stress sensor, plays a critical role in *E. coli*-induced LPM inflammasome activation. Surprisingly, pretreatment of NLRP3 inhibitor, MCC950, had no effects on IL-1β release from *E. coli*-infected control and *Pacc1*^−/−^ LPMs (Fig. S5 I), suggesting other mechanisms might play a more important role, such as flagellin-NAIP-NLRC4 or non-canonical LPS-CASP11 pathways. Taken together, these results suggest that the phagosome bacterial degradation promotes inflammasome activation and pyroptosis likely through the release of bacterial ligands from phagosomes in LPMs.

Previous studies have shown that reconstituted gasdermin D-NT protein forms large pores on liposomes containing cardiolipin, a phospholipid found in bacterial membranes (Ding et al., 2016; Liu et al., 2016). Moreover, co-culturing *E. coli* with purified gasdermin D-NT or supernatants from inflammasome-activated cells kills bacteria, suggesting a direct bactericidal activity (Liu et al., 2016). To test if the augmented pyroptosis in PAC-deficient LPMs directly facilitate peritoneal bacterial killing, we first examined the release of gasdermin D by LPMs upon *E. coli* infection. Consistent with increased inflammasome activation, *Pacc1*^−/−^ LPMs secreted more gasdermin D to the supernatants than control cells (Fig. S5 J). Importantly, treating fresh *E. coli* culture with antibiotic-free supernatants from infected *Pacc1*^−/−^ LPMs more strongly inhibited bacterial growth than antibiotic-free supernatants from infected control LPMs. To further validate the role of the released gasdermin D, we also treated *E. coli* with antibiotic-free supernatants from infected *Gsdmd^−/−^* LPMs, which showed no inhibition in bacterial growth (Fig. 4 K and Fig. S5 K). While contributions from other secreted factors cannot be ruled out, these findings suggest that extracellular gasdermin D could contribute to bacterial growth inhibition *in vitro*. To test this *in vivo*, we collected peritoneal fluids from mice after peritoneal *E. coli* infection and observed a higher gasdermin D release in *Cx3cr1*-PAC^−/−^ mice than controls (Fig. 4 L). Interestingly, we also observed more surface gasdermin D puncta together with propidium iodide (PI) signals in bacteria isolated from the peritoneum of *Cx3cr1*-PAC^−/−^ mice (Fig. 4 M), further supporting gasdermin D pore-forming as a potential bactericidal mechanism.

### LPM depletion-adoptive transfer model reveals a role for gasdermin D in bacterial clearance

We next sought to further examine the role of LPM inflammasome activation and pyroptosis in bacterial clearance by focusing on their main effector gasdermin D. We developed a novel depletion-adoptive transfer model for LPMs (Fig. 4 N and Fig. S5 M). Clodronate-containing liposomes can effectively deplete cavity resident macrophages, such as peritoneal and pleural macrophages (van Rooijen and Sanders, 1997). Indeed, peritoneal injection of clodronate liposomes specifically depleted LPMs without affecting SPMs (Fig. S5 L). To further minimize the depletion-induced inflammation, we waited 12 days before the adoptive transfer. We examined the peritoneal cell composition post-depletion, revealing minimal neutrophil recruitment, indicating low inflammation, and modest monocyte recruitment, likely to replenish depleted LPMs (Fig. S5 N). Using CellTrace dyes, we labeled MACS-isolated LPMs from donor mice to monitor their viability post-adoptive transfer. FACS analysis confirmed clear gating and high viability of transferred cells (Fig. S5 O). Notably, adoptive transfer of WT control LPMs significantly protected mice from lethality due to peritoneal *E. coli* infection (Fig. S5 P). While recruited monocytes may contribute to this protection, the phagocytic nature of transferred LPMs likely plays a primary role. With this new tool, we first tested if PAC truly plays a LPM cell-autonomous role in bacterial clearance. To this end, we transferred *Pacc1*^−/−^ LPMs into LPM-depleted mice, focusing on CellTrace-labeled LPMs and their bactericidal activities. Consistently, we observed an augmented anti-bacterial activity of the KO LPMs compared to control LPMs after peritoneal *E. coli* infection. This included a decrease in intracellular *E. coli*-GFP signals in LPMs (Fig. 4 O), indicating an increased bactericidal activity, and a reduction in peritoneal *E. coli* number (Fig. 4 P), suggesting a better infection resolution. These data corroborate the findings based on *Cx3cr1*-PAC^−/−^ and *LysM*-PAC^−/−^ mice, supporting a role in bacterial clearance for the PAC channel specifically in LPMs. They further validate the depletion-adoptive transfer model as a reliable method to investigate the function of LPMs.

To examine the role of LPM pyroptosis in bacterial clearance, we next adoptively transferred LPMs isolated from *Gsdmd^−/−^* mice to LPM-depleted mice and then performed peritoneal infection with *E. coli*-GFP (Fig. 4 N). As expected, *Gsdmd* deletion protected the transferred LPMs from pyroptotic cell death during infection (Fig. 4 Q). Intracellular GFP signals were similar between transferred control and *Gsdmd^−/−^* LPMs, suggesting that gasdermin D in LPMs does not affect *E. coli* uptake or killing (Fig. 4 R). However, interestingly, we observed an increase in *E. coli* number within peritoneal fluids in mice transferred with *Gsdmd^−/−^* LPMs (Fig. 4 S). This data indicates that gasdermin D in LPMs is involved in promoting extracellular bacterial clearance, consistent with a direct bacterial killing activity previously proposed (Ding et al., 2016; Liu et al., 2016) and further supported by our findings (Fig. 4 K-S and Fig. S5 J and K). Together, these results suggest that enhanced phagosome bacterial degradation in PAC-deficient LPMs promotes inflammasome activation and gasdermin D release, likely contributing to improved bacterial clearance and resolution of *E. coli* infection.

### PAC deletion in macrophages mitigates peritoneal inflammation and promotes resolution of peritoneal *E. coli* infection

The balance between cellular activation for anti-microbial activities and subsequent inflammation are crucial determinants for host survival during infections (Guo et al., 2015a). The hyperactivation of IFN and inflammasomes in PAC-deficient LPMs prompted us to examine the overall peritoneal inflammation status in the KO mice. To achieve that, we first performed a comprehensive secretome analysis with peritoneal fluids after peritoneal *E. coli* infection (Fig. S4 D). Surprisingly, we observed a consistent reduction in several inflammatory cytokines and chemokines in both *Cx3cr1*-PAC^−/−^ and *LysM*-PAC^−/−^ mice compared to controls, including IL-6, macrophage migration inhibitory factor (MIF), and CCL8 (Fig. 5 A and B). Interestingly, CXCL2 and IL-1β levels were decreased in *LysM*-PAC^−/−^ mice, contrasting with *in vitro* LPM infection results (Fig. S4 H). This difference suggests the contribution of other peritoneal cells to the cytokine secretion and highlights bacteria as the primary driver of peritoneal inflammation. MIF is a major driver for peritoneal inflammation in a peritonitis-induced sepsis model (Tilstam et al., 2021). IL-6 and IL-1β are key proinflammatory cytokines, while CCL8 and CXCL2 as major chemoattractants for neutrophils and monocytes. Only one chemokine, CXCL9, was increased in *Cx3cr1*-PAC^−/−^ mice. CXCL9 has been shown to protect mice against colonic bacterial infection although the underlying mechanism is unclear (Reid-Yu et al., 2015). We also observed a very modest increase of IL-33 in *LysM*-PAC^−/−^ mice. Given its very low concentration, the significance of such an increase remains unclear. Overall, the secretome analysis revealed a less inflammatory environment in the peritoneum of macrophage-specific PAC KO mice after *E. coli* infection. These results suggest that although PAC-deficient LPMs were hyperactivated upon bacterial infection, enhanced bacterial clearance may dampen overall peritoneal inflammation due to reduced activation of other peritoneal cells, such as SPMs as well as recruited proinflammatory neutrophils and monocytes.

**Figure 5.**
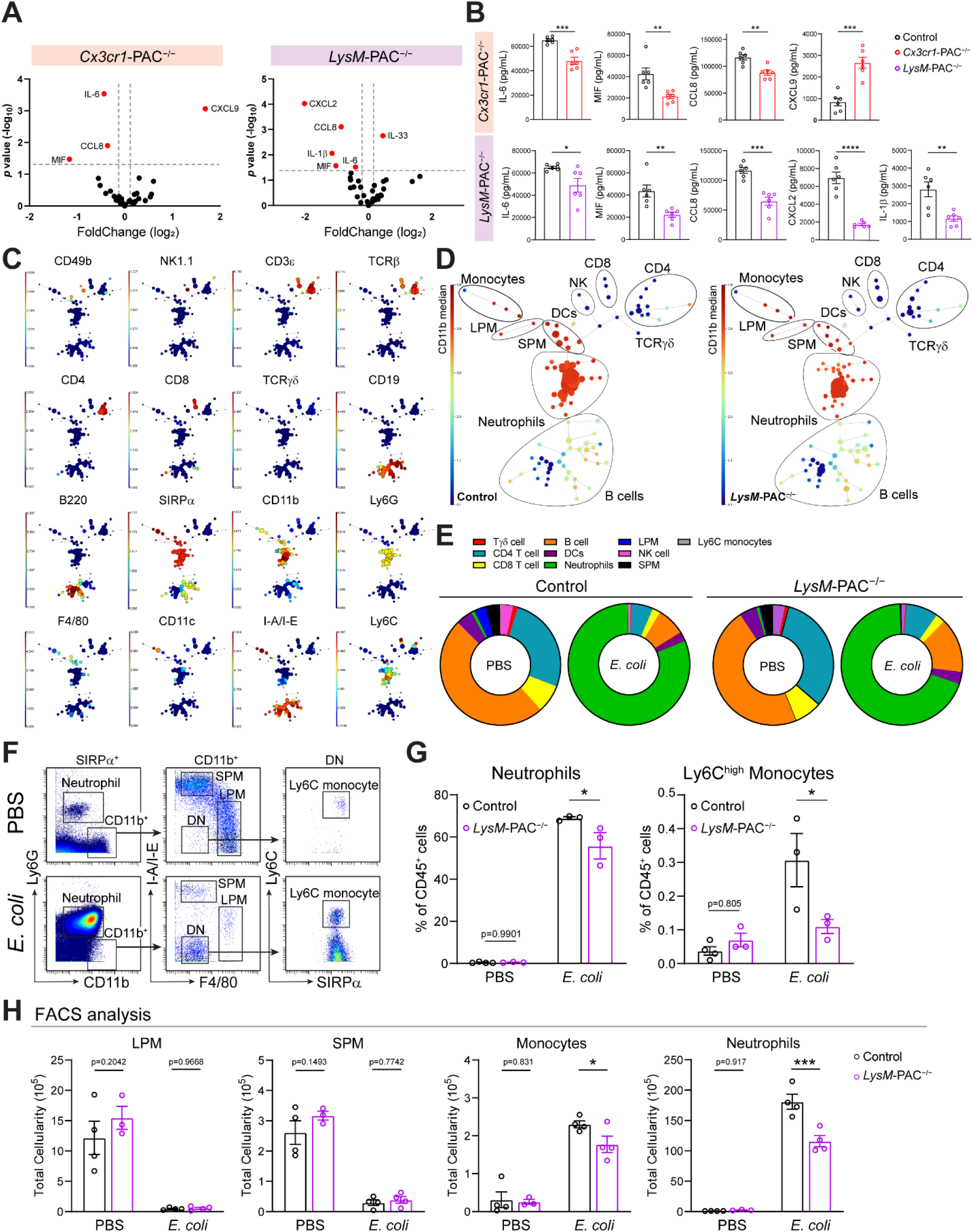
PAC deletion in macrophages mitigates peritoneal inflammation after peritoneal *E. coli* infection. **A,** Volcano plot of ELISA assays from peritoneal fluids 4 hours post peritoneal infection. Highlighted proteins represent top differentially secreted proteins with p < 0.05 and average log-transformed fold-change > 0.25. **B**, Quantification of highlighted proteins in (**A**), with a minimum concentration above 100 pg/mL. Data represented from n = 6 mice. **C**, Feature plot generated by FlowSOM analysis for cyTOF dataset with a minimum population size of 2%. Red nodes indicate an abundance in expression for indicated cell marker. Each trajectory suggests a lineage of cell type. **D**, FlowSOM analyzed colored tree for immune cell types in cyTOF dataset. **E**, Donut plots of peritoneal immune cell types within cyTOF dataset. **F**, Manual gating strategy for the cyTOF dataset. DN, double negative. **G**, cyTOF quantification of neutrophil and Ly6C^high^ monocyte populations. Data represented from n > 3 mice. **H**, FACS quantification of indicated cell types in peritoneal fluids. The PBS control data corresponds to Fig. S2C. Data represented from n > 3 mice. All control groups represent *Pacc1^F/F^* genotype. Data are reported as mean ± SEM between independent experiments. Unpaired t-test for **B**. Two-way ANOVA with Sidak’s test for **G, H**. *p < 0.05, **p < 0.01, ***p < 0.001, ****p < 0.0001.

To evaluate the contribution of infiltrated immune cells to peritoneal inflammation, we employed an unbiased high-dimensional mass cytometry (termed cytometry by Time-Of-Flight, cyTOF) analysis to examine changes in immune cell populations upon peritoneal infection. Given the chemokine receptor nature of CX3CR1, we focused our analysis on *LysM*-PAC^−/−^ instead of *Cx3cr1*-PAC^−/−^ mice. We isolated cells from the peritoneal lavage after *E. coli* infection and subjected them to cyTOF analysis. We first unmasked major peritoneal cell populations using algorithmic clustering, which discerned nine major immune populations, including LPMs, SPMs, neutrophils, dendritic cells, monocytes, mast cells, NK cells, T cells (CD4, CD8, and γδT cells), and B cells (Fig. 5 C and D). As expected, *E. coli* infection in control mice resulted in dramatic increases in neutrophils and monocytes, the two major infiltrated immune cells during peritoneal bacterial infection. Consistent with a less inflammatory environment, the recruitment of neutrophils as well as highly proliferative Ly6C^high^ monocytes was markedly reduced in *LysM*-PAC^−/−^ mice compared to controls (Fig. 5 E-G). The decreased recruitment of these infiltrating cells was also validated with FACS analysis (Fig. 5 H). Together with the secretome analysis, these results suggest that enhanced bacterial clearance by PAC-deficient LPMs reduces the subsequent recruitment of proinflammatory cells, thereby further mitigating peritoneal inflammation.

To further evaluate the role of PAC in infection resolution *in vivo*, we conducted a time-course analysis of bacterial load in peritoneal fluids after peritoneal *E. coli* infection. Consistent with the enhanced bactericidal activity in LPMs, both *Cx3cr1*-PAC^−/−^ and *LysM*-PAC^−/−^ mice exhibited a reduced peritoneal bacterial load compared to controls (Fig. 6 A). Uncontrolled peritoneal bacterial infection can lead to systematic sepsis and even death (Brun-Buisson et al., 1995). In line with the decreased bacterial load and dampened peritoneal inflammation, both macrophage-specific PAC KO mice showed alleviated bacteremia in the tail blood and were largely protected from sepsis-induced mortality (Fig. 6 B and C). In addition to WT *E. coli* strain, we also observed enhanced bacterial clearance and improved survival for peritoneal infection of a clinically relevant carbapenem-resistant strain of *E. coli* (cpm-resistant *E. coli*) (Fig. 6 D and E). These data demonstrate that enhanced phagosome bacterial degradation together with downstream IFN and inflammasome activation in PAC-deficient macrophages promotes disease resolution and reduces mortality against peritoneal *E. coli* infections *in vivo*.

**Figure 6.**
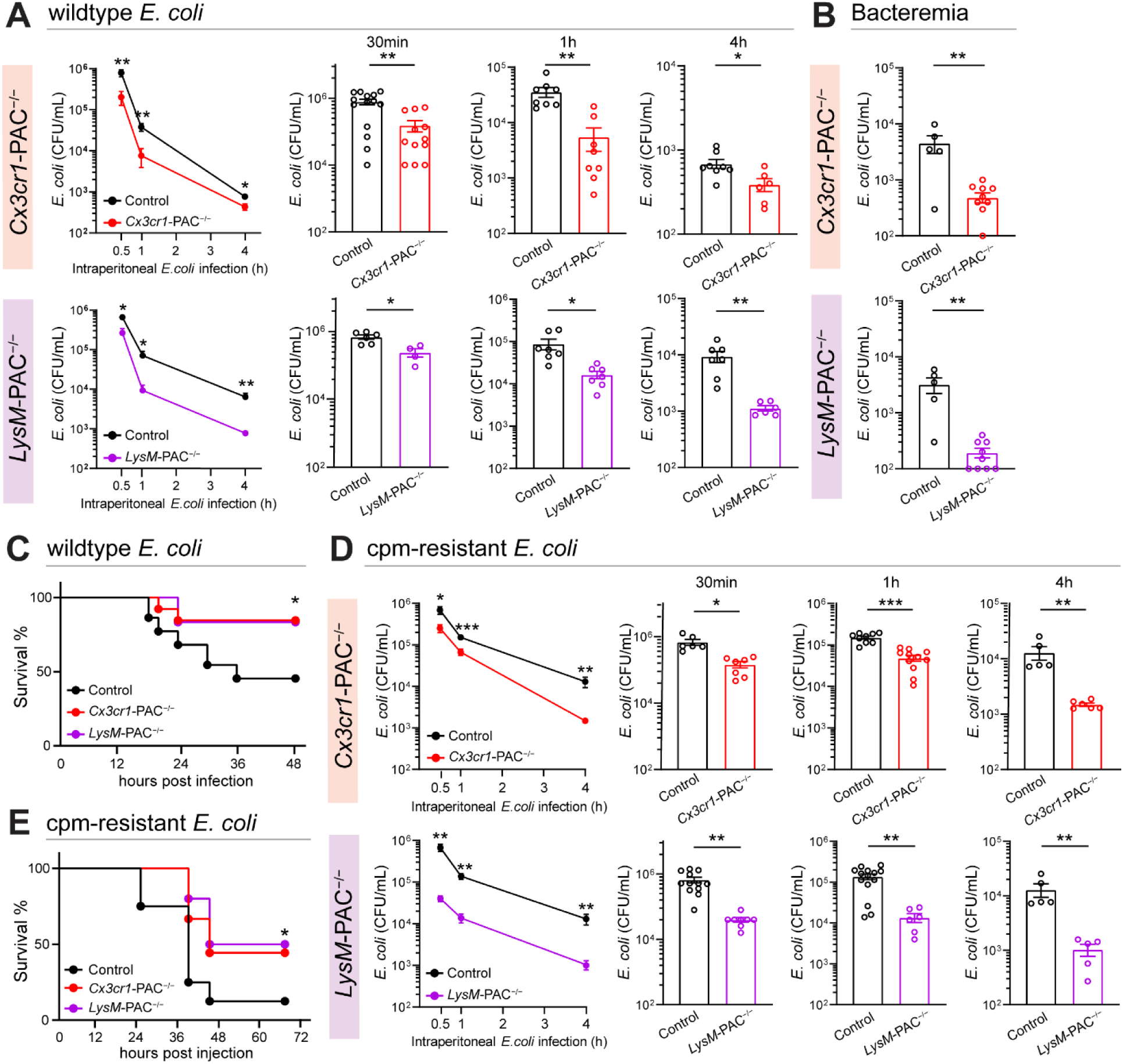
PAC deletion in macrophages promotes resolution of peritoneal *E. coli* infection. **A**, Bacterial culture from peritoneal fluids after *E. coli* peritoneal infection. Data represented from n > 4 mice. **B**, Bacterial culture from tail blood after *E. coli* peritoneal infection. Data represented from n > 5 mice. **C**, Survival of mice after *E. coli* peritoneal infection at 2 × 10^7^ CFU/mL; WT n = 19. *Cx3cr1*-PAC^−/−^ n = 13. *LysM*-PAC^−/−^ n = 12. **D**, Bacterial culture from peritoneal fluids after cpm-resistant *E. coli* peritoneal infection. Data represented from n > 4 mice. **E**, Survival of mice after cpm-resistant *E. coli* peritoneal infection at 10^8^ CFU/mL; WT n = 8. *Cx3cr1*-PAC^−/−^ n = 9. *LysM*-PAC^−/−^ n = 10. All control groups represent *Pacc1^F/F^* genotype. Data are reported as mean ± SEM between independent experiments. Unpaired t-test for **A, B, D**. Mantel-Cox test and Gehan-Breslow-Wilcoxon test for **C**, **E**. *p < 0.05, **p < 0.01, ***p < 0.001.

## Discussion

Our study identifies that PAC, a newly identified chloride channel, as a critical negative regulator for phagosome maturation and anti-bacterial immunity. PAC-deficient macrophages exhibited increased phagosomal acidification and protease activation, leading to enhanced bactericidal activity and improved mouse survival upon peritoneal *E. coli* infection (Fig. 7). These findings not only highlight the importance of the PAC channel in phagosome maturation and bacterial clearance but also provide an ideal model to examine the role of phagosome-mediated degradation in shaping host defense responses against bacterial infection.

**Figure 7.**
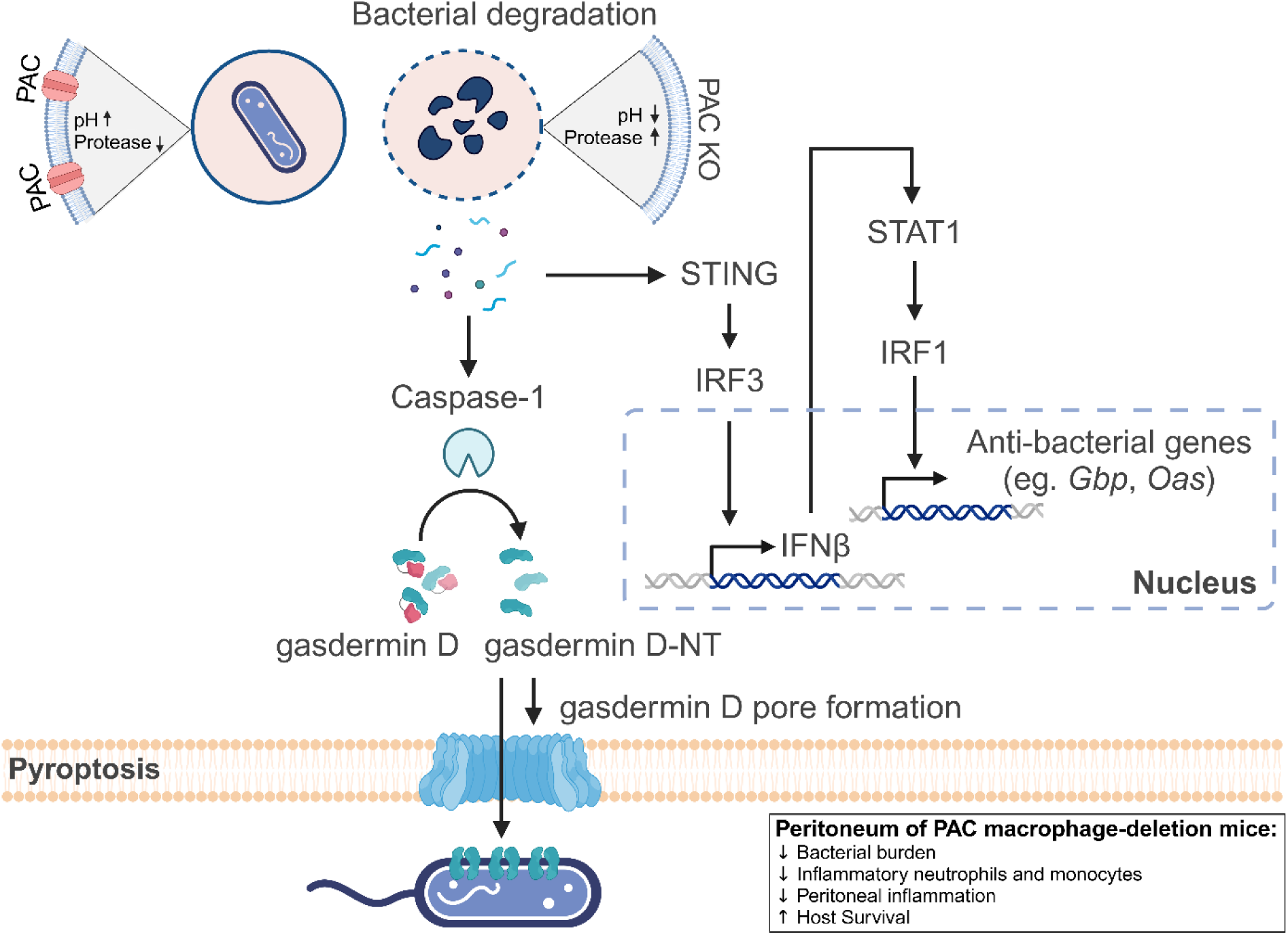
Summary diagram. PAC regulates phagosome maturation and phagosome-mediated antibacterial pathways in peritoneal macrophages

We find that enhanced phagosome bacterial degradation in PAC-deficient LPMs boosted inflammasome activation. Inflammasomes, activated by diverse sensors, respond to pathogenic ligands such as bacterial LPS, flagellin, DNA, and RNA. For instance, NAIP5-NLRC4 detects flagellin, AIM2 senses bacterial DNA, and NLRP3 recognizes bacterial toxins (Barnett et al., 2023; Paidimuddala et al., 2023). As multiple bacterial ligands may be released into the cytoplasm following phagosomal degradation, the observed enhanced inflammasome activation likely results from the combined activity of multiple sensors. Additionally, canonical pathways, such as ATP release from dying neighboring cells, may also contribute to this activation. This may also explain the residual IL-1β secretion after E64d treatment in both control and *Pacc1*^−/−^ LPMs (Fig. 4 G). Further studies are needed to delineate the relative contributions of various pathways.

Augmented phagosome bacterial degradation in PAC-deficient LPMs also stimulated STING-IRF3, another key innate immune pathway, leading to IFNβ production. Interestingly, a recent preprint reported that evolutionarily diverse bacteria, including *E. coli*, stimulate IFN responses in macrophages after phagosome-mediated bacteriolysis in a membrane bound-TLR-independent, but STING-dependent manner. The authors further provided evidence that two CDN transporters LRRC8A (as known as SWELL1) and SLC46A2 are responsible for the phagosome-mediated cytoplasmic activation of STING (Stephanie A. Ragland, 2022). STING and various other cytoplasmic PRRs are traditionally thought to be activated by intracellular pathogens, such as viruses and bacteria escaping phagosomes, or the release of mitochondrial DNA to the cytoplasm during infections (Abdullah and Knolle, 2014; Man et al., 2016; West et al., 2015). However, our findings and the new study highlight the engagement of phagosome-mediated bacteriolysis in innate immune activation by activating cytoplasmic PRRs with the released bacterial ligands (CDNs, flagellin, and nucleic acids) from phagosomes. Beyond phagosome-mediated release, mitochondrial DNA release could be another phagosome-independent STING activator. Future research will determine if and how various STING-activating pathways together contribute to the IFN responses in different immune cells upon infection.

Besides the conventional phagocytic bacterial killing, our comprehensive analysis on PAC-deficient LPMs in the peritoneal *E. coli* infection model reveals two anti-bacterial mechanisms that are downstream of elevated phagosome degradation. First, the activation of STING-IRF3 signaling pathway drives the production of IFNβ, which upregulates the expression of key anti-microbial factors, including GBP and OAS family proteins (Fisch et al., 2023; Hornung et al., 2014; Leisching et al., 2019; Man et al., 2016). Second, enhanced inflammasome activation and pyroptosis in LPMs further facilitate bacterial clearance by releasing gasdermin D to the peritoneum, which forms pores on the bacterial surface (Ding et al., 2016; Liu et al., 2016). Additional mechanisms of bacterial removal may also exist. For example, the pyroptotic LPMs may form bacterial traps from cell corpses, which eventually promote efferocytosis by other phagocytes (Jorgensen et al., 2016).

The commonly used *Cx3cr1*-Cre and *LysM*-Cre models mediate gene deletion in all macrophages, including SPMs, monocytes, and potentially other cell types (Clausen et al., 1999; Yona et al., 2013). To circumvent this limitation, we developed a depletion-adoptive transfer model for LPMs, complementing our conditional KO approach and validating PAC’s specific role in LPM-mediated bacterial clearance. It also revealed the involvement of LPM inflammasome activation in peritoneal bacterial killing. Notably, the peritoneal environment may not fully recover to normal after 12 days of clodronate liposome treatment, as indicated by the recruited monocyte population. Nevertheless, we expect that this depletion-adoptive transfer model will be useful for the field to investigate the biology of LPMs with proper controls.

Our RNA-seq analysis of *E. coli*-infected LPMs revealed that the upregulated genes were mainly associated with signatures of bacterial sensing and killing instead of conventional inflammation pathways. This observation aligns with the notion that LPMs serve as the dedicated phagocytes, while SPMs and the recruited neutrophils and monocytes are major drivers for peritoneal inflammation during bacterial infections (Louwe et al., 2021; Teh et al., 2022; Tilstam et al., 2021; Vega-Perez et al., 2021). The expression of these anti-bacterial molecular signatures, but not proinflammatory genes (i.e. *Nos2*, *Tnf* or *Il6*) were further increased in PAC-deficient LPMs when phagosome-mediated bacterial degradation was enhanced. This is consistent with the finding that *Cx3cr1*-PAC^−/−^ and *LysM*-PAC^−/−^ mice not only exhibited a better LPM-mediated bacterial clearance during disease resolution, but also showed mitigated peritoneal inflammation. These results further suggest that the peritoneal bacterial load is the major determinant for peritoneal inflammation, likely by regulating the activation of SPMs and other infiltrated immune cells.

The negative role of PAC in phagosome acidification in macrophages is consistent with a model previously proposed for endosomes that PAC is activated by acidic pH and releases the counterion Cl^−^ from organelle lumen, thereby preventing further acidification (Osei-Owusu et al., 2021). The increase in phagosomal protease activities in PAC-deficient macrophages is likely due to more efficient phagosomal acidification. Loss of the PAC channel may also lead to a higher Cl^−^ concentration, which could elevate protease activities as well (Wu et al., 2023). In addition to proteases, the activity of other hydrolytic enzymes in phagosomes, such as glycosidases, lipases, and nucleases, is likely also enhanced in PAC-deficient macrophages. LPS treatment typically upregulates gene expression. However, PAC expression was downregulated by LPS, which in turn is required for LPS-induced phagosome activation. This provides an elegant mechanism for macrophages to facilitate phagosome maturation in response to bacterial infections, while preserving the housekeeping function of PAC in maintaining endosomal pH homeostasis under normal physiological conditions (Osei-Owusu et al., 2021; Zeziulia et al., 2022). Although macrophages are efficient in killing most bacteria, some pathogens, such as *Mycobacterium* and *Legionella pneumophilia*, can survive and adapt after phagocytosis (Horwitz and Maxfield, 1984; Queval et al., 2017). They achieve this mainly by hijacking the host acidification machinery and preventing phagosomal acidification. In the future, it will be interesting to test if the effector proteins from these bacteria can directly target and, perhaps, activate the PAC channel to interfere with acidification.

PAC deletion presents beneficial effects in enhancing macrophage bactericidal activities against both WT and cpm-resistant *E. coli* infections, leading to reduced lethality in mice. By promoting phagosome-mediated bacterial degradation, the effect of PAC deletion likely extends beyond *E. coli*, to other clinically relevant bacteria, especially those sensitive to phagosome degradation, such as *Streptococcus pneumoniae* (Ip et al., 2010). Our study thus suggests the PAC channel as a potential therapeutical target for the treatment of bacterial infectious diseases. Due to the lack of molecular identity until recently, there is no potent and selective PAC inhibitor available. We previously developed a high-throughput cell-based fluorescence quenching assay for the PAC channel and applied it to successfully identify its underlying gene. This assay will offer a promising avenue for screening the PAC inhibitors in the future, which can be tested for their potential in combating bacterial infections.

## Acknowledgments

We thank Dr. Andrea Cox for providing human THP1 cells, Dr. Won Jin Ho and Johns Hopkins Sidney Kimmel Comprehensive Cancer Center (SKCCC) Flow/Mass Cytometry Core for the assistance on cyTOF experiments, as well as Dr. Nathan Archer and members of the Qiu lab for valuable discussion.

## Funding

National Institutes of Health grant R35GM124824 (ZQ)

National Institutes of Health grant R01NS118014 (ZQ)

National Institutes of Health grant RF1NS134549 (ZQ)

McKnight Scholar award (ZQ)

Klingenstein-Simon Scholar award (ZQ)

Sloan Research Fellowship in Neuroscience (ZQ)

Karen Toffler Scholar Program (YY)

## Author contributions

Conceptualization: HYC, JChu, ZQ

Methodology: HYC, JChu, JChen, YY, SY, NL, KHC, XD

Investigation: HYC., JChu, JChen

Visualization: HYC

Funding acquisition: ZQ

Project acquisition: HYC, JChu, ZQ

Supervision: ZQ

Writing – original draft: HYC, ZQ

Writing – review & editing: HYC, ZQ

## Materials and Methods

### Contact for reagent and resource sharing

Further information and requests for resources should be directed to and will be fulfilled by the Lead Contact, Zhaozhu Qiu.

### Material availability

PAC KO THP-1 (gRNA1 and gRNA2), PAC overexpressing Raw264.7, and PAC-mCherry expressing Raw264.7 cells were generated in this study. *Cx3cr1*-PAC^−/−^, *Cx3cr1*-SWELL1^−/−^, and *LysM*-PAC^−/−^ mice were generated in this study. All will be made available from the lead contact upon request.

### Experimental Model and Subject Details

#### Mice

The use of all mice in the study was approved by the Johns Hopkins University Animal Care and Use Committee, under approved protocol MO22M17. All mice were housed in a specific pathogen free (SPF) environment, under standard 12-hour light/12-hour dark cycle with *ad libitum* access to food and water. PAC*^F/F^* mice were generated in the laboratory as previously described(Yang et al., 2019), while *Gsdmd*^−/−^ (C57BL/6N-*Gsdmd^em4Fcw^*/J), *Cx3cr1*-Cre [B6J.B6N(Cg)-*Cx3cr1^tm1.1(cre)Jung^*/J] and *LysM*-Cre [B6.129P2-*Lyz2^tm1(cre)Ifo^*/J] were purchased from The Jackson Laboratory (strain #032410, #025524 and #004781, respectively). *Cx3cr1*-PAC^−/−^ and *LysM*-PAC^−/−^ mice were generated by crossing *Cx3cr1*-Cre or *LysM*-Cre mice with PAC*^F/F^* mice, with *Cx3cr1*^Cre/WT^;PAC*^F/F^* and *LysM* ^Cre/WT^;PAC*^F/F^* considered as *Cx3cr1*-PAC^−/−^ and *LysM*-PAC^−/−^, respectively. Adult male or female mice between 8-12 weeks of age were used for experiments as specified with age- and sex-matched littermates as controls.

#### Bacterial Culture

Wildtype (WT) *Escherichia coli* (*E. coli*), GFP-expressing *E. coli*, and cpm-resistant *E. coli* strains were purchased from American Type Culture Collection (ATCC, #25922, #25922GFP, and #BAA-2469 respectively). All handling of the infectious bacterial strains strictly adhered to instructions provided by the Johns Hopkins Biosafety Office. The use of bacterial strains in the study was approved by Johns Hopkins Institutional Biosafety Committee (IBC).

Bacterial cultures were prepared following the instructions provided by ATCC. Briefly, wildtype *E. coli* was grown in LB broth. *E. coli*-GFP was grown in LB broth supplemented with carbamacilin (100 μg/mL). Cpm-resistant *E. coli* was grown in Trypticase Soy Broth (TSB) supplemented with Imipenem (25 μg/mL). All cultures were incubated aerobically at 37°C overnight. Subsequently, bacteria were subcultured (1:50) in respective fresh media for 4 hours for the determination of colony formation units (CFU) that were used for the subsequent infection experiments.

#### Cell Culture

Details of BMDM culture have been previously reported (Zeziulia et al., 2022). Briefly, BMDMs were freshly isolated and cultured from adult C57BL6/J mice (8-12 weeks old). Bone marrow was flushed from both femurs and tibiae with a 26-gauge needle syringe into a 50 mL conical tube. Hematopoietic stem cells were collected through 300xg 10 min of centrifugation at 4°C. BMDMs were differentiated by culturing stem cells in 10 cm petri dishes in L929-differentiation medium or full RPMI 1640 medium (10% HI-FBS, 2mM L-glutamine, 10mM MEM non-essential amino acids, 100 units/mL penicillin, and 100 mg/mL streptomycin) supplemented with mouse recombinant M-CSF (40 ng/mL) for 6 days. L929-differentiation medium was made from 70% RPMI 1640 medium (20% heat-inactivated fetal bovine serum (HI-FBS), 100 units/mL penicillin, and 100 mg/mL streptomycin) supplemented with 30% L929 mouse fibroblast-conditioned medium. During the differentiation, 2-3 mL of freshly made respective medium was supplemented every 2 days. On day 6, matured BMDMs were then subcultured using macrophage detachment buffer (10mM EDTA in PBS) for following experiments.

Raw264.7 mouse macrophage cell line was purchased through ATCC (TIB-71) and cultured following manufacturer’s instructions. Briefly, cells were maintained in Dulbecco’s modified Eagle’s medium (DMEM) supplemented with 10% HI-FBS, 100 units/mL penicillin, and 100 mg/mL streptomycin at 37°C in humidified 95% CO_2_ incubator. THP-1 human monocytes, generously provided by Dr. Andrea Cox (Johns Hopkins University), were cultured in full RPMI 1640 medium at 37°C in a humidified 95% CO_2_ incubator. THP-1 macrophages were differentiated by treating the cells with 5 ng/mL of PMA for 24 hours, followed by an additional 24-hour resting in RPMI 1640 medium before subsequent experiments.

### Method Details

#### Genotyping and Characterization of Mice

The extraction of genomic DNA from tail samples was performed with Extracta DNA Prep kit (Quanta Bio). Samples were digested with 20 μL of digestion buffer for 30 min at 95°C, and stopped with 20 μL of stopping buffer. 1 μL of genomic DNA sample was further used as template for PCR using AccuStart II GelTrack PCR SuperMix (Quanta Bio). PCR products were electrophoretically separated with a 2-3% agarose gel and visualized with SYBR safe DNA gel stain (Thermo Fisher Scientific). Genotype was determined using the following primers:

*Cx3cr1*-Cre:

Common primer: 5’-GCA GGG AAA TCT GAT GCA AG-3’
Mutant primer: 5’-GAC ATT TGC CTT GCT GGA C-3’
Wildtype primer: 5’-CCT CAG TGT GAC GGA GAC AG-3’

*LysM*-Cre:

Common primer: 5’-AAG GAG GGA CTT GGA GGA TG-3’
Mutant primer: 5’-ACC GGT AAT GCA GGC AAA T-3’
Wildtype primer: 5’-GTC ACT CAC TGC TCC CCT GT-3’

*Gsdmd*^−/−^:

Forward primer: 5’-ACA CCC CTG AAC CCA TAT CC-3’
Reverse primer: 5’-GCA CTC ACC CCA TTT AGA GC-3’

*Casp1*^−/−^:

Common primer: 5’-ATG GCA CAC CAC AGA TAT CGG-3’
Mutant primer: 5’-GAG ACA TAT AAG GGA GAA GGG-3’
Wildtype primer: 5’-TGC TAA AGC GCA TGC TCC AGA CTG-3’

LoxP sites at *Pacc1* exon 2:

Forward primer: 5’-GAA GCC AGG CCA TTC TTT TT-3’
Reverse primer: 5’-GCT CAA GGA AAC CAC TGA GG-3’

LoxP sites at *Swell1:*

Forward primer: 5’-GAA ACA GTG AGG GTT CAG GAG-3’
Reverse primer: 5’-ATG TTG TTG GGA GAC AGA TAC C-3’

#### Peritoneal Bacterial Infection

The CFU of each bacterium freshly grown after 4 hours was determined by LB plate culture. For the peritoneal infection, bacterium was resuspended in sterile PBS. Mice were peritoneally infected by the injection of 200 μL of bacterial solution with indicated CFU and monitored daily for health and survival following institutional guidance.

#### Peritoneal Lavage

Mice were euthanized before the procedure. Peritoneal lavage was carried out by injecting 7 mL of sterile PBS into the peritoneal cavity, followed by gentle abdominal massage for homogenization. Afterward, a skin incision was made and a 26-guage syringe was used to collect peritoneal fluid. 5 mL of the injected PBS from each mouse was collected and transferred into a 15 mL conical tube. Centrifugation at 300xg for 5-10 minutes at 4°C was performed to isolate peritoneal cells or obtain peritoneal fluid for subsequent analyses. ACK lysis buffer (Quality Biological) was used for the removal of red blood cells. For peritoneal fluid secretome analyses, 1 mL of PBS was injected, and 0.6-7 mL of the injected fluid was subsequently harvested.

#### Flow Cytometry (FACS) Analysis

Single cell suspension was prepared from peritoneal lavage or cell culture and resuspend in FACS buffer (2% FBS, 2 mM EDTA in sterile PBS). Live versus Dead cells were stained in PBS using Live/Dead Fixable Aqua Dead Cell Stain (Thermo Fisher Scientific). Fc receptors were then blocked by incubation with TrueStain FcX anti-CD16/32 antibody (BioLegend) at 4°C for 20 minutes, and the cells were subsequently stained with fluorophore-conjugated antibodies against the mouse antigens listed in the Supplemental Material Table, for 20 minutes at 4°C. For primary antibodies without fluorophore, subsequent fluorophore-conjugated secondary antibodies were used with the same staining approach. For the time-traced bacterial clearance analysis, the mean fluorescence intensity (MFI) of GFP within LPM or neutrophils was acquired following specified infection time. FACS analysis of intracellular GBP5 was carried out using Cyto-Fast Fix/Perm Buffer set (BioLegend), following manufacturer’s instructions. Detection of the cell death was performed, after cell surface labeling, using Annexin V and 7-AAD staining (BioLegend), following manufacturer’s instructions. Data was acquired on a CytoFLEX LX (Beckman Coulter) and analyzed using FlowJo (BD).

#### Mouse LPM depletion-adoptive transfer model

Male WT mice of 8-12 weeks old were subjected to LPM depletion with peritoneal injection of 100 uL PBS-loaded or clodronate-loaded liposomes (Liposoma). 12 days post depletion, LPMs isolated from WT, *Pacc1*^−/−^, or *Gsdmd*^−/−^ mice were counted, labeled with CellTrace CSFE or CTV dyes (Invitrogen), and adoptively transferred to the peritoneum of LPM-depleted mice. After 24 hours of transfer, mice were peritoneally infected with *E.coli*-GFP to evaluate LPM bactericidal activities and disease resolution. Otherwise, 12 days post depletion or 24 hours after LPM transfer, mouse peritoneal cells were acquired by peritoneal lavage for the FACS analysis determination of cell composition.

#### Secretome ELISA Analysis

Peritoneal fluid prepared from peritoneal lavage, or LPM culture medium post infection was subjected to LegendPlex multi-analyte ELISA assay (BioLegend) or individual ELISAs following manufacturer’s instruction. For LPM supernatants, freshly isolated LPMs were plated at a density of 5 × 10^5^ cells/well in 12-well plates. For *E. coli* infection, bacterium at MOI 20 was simultaneously plated in 12-well plates. Both peritoneal fluids and cell supernatants were collected, centrifugated at 12000xg to remove dead cells and debris before subjected to ELISA assays. For multiplex ELISA assays, samples were prepared with multiple dilution factors to ensure cytokines fell within their dynamic detection ranges. Specifically, pre-defined LegendPlex panels of Mouse Cytokine Release Syndrome Panel, Mouse Cytokine Panel 2, and Mouse Proinflammatory Chemokine Panel 2 were selected for analyses. Additionally, IL-18 duoset ELISA (R&D Systems), IFNβ duoset ELISA (R&D Systems), MIF Legend MAX ELISA (BioLegend), and CXCL1 duoset ELISA (R&D Systems) were utilized for the purpose of comprehensiveness. For the gasdermin D release quantification, Mouse gasdermin D ELISA Kit (Abcam) was used to determine gasdermin D release in LPM supernatant following manufacturer’s instruction.

#### LPM Purification and Isolation

LPMs were isolated from peritoneal fluids by negative selection after immunomagnetic depletion of B cells, SPMs, T cells, and neutrophils with peritoneum mouse macrophage isolation kit (Miltenyi Biotec). After immunomagnetic depletion, LPM preparations had a purity > 90%. For immunoblotting analysis, isolated LPM was immediately lysed in cell lysis buffer. For immunofluorescence analysis, isolated LPM was cultured on coverslips in full RPMI 1640 medium at 37°C for an additional 20 min for adhesion before fixation. For secretome analysis, isolated LPM was immediately infected with *E. coli* in full RPMI 1640 medium at 37°C for 24 hours before supernatant collection.

#### Gene Transduction and CRISPR Deletion

PAC KO THP-1 cells were generated by CRISPR-Cas9 technology. Guide RNA1 (GGACCGAGAAGACGTTCTTC) and guide RNA 2 (GTGGATCGCTATGATGCCCC) targeting exon 3 of human *PACC1*/*TMEM206* (PAC) was cloned into LentiCRISPR-v2-PuroR plasmid, which was a gift from Dr. F. Zhang (Addgene plasmid #98290). Lentiviral particles containing PAC sgRNA1 or sgRNA2 were packaged using the third-generation lentiviral system and used to infect THP-1 cells. 24 hours after infection, the medium was changed to fresh RPMI culture medium. Cells were then treated with puromycin (1 μg/ml) for 3-5 days to select for successfully transduced cells. PAC protein expression was validated in puromycin-resistant cells using protein immunoblotting or whole-cell recording for PAC-specific currents.

PAC overexpressing and mCherry-tagged PAC overexpressing Raw264.7 cells were generated using plasmids as previously described (Osei-Owusu et al., 2021) and packaged into lentiviral particles using the third-generation lentiviral system. 24 hours after infection, the medium was changed to fresh DMEM culture medium. Cells were then FACS sorted for mCherry-positive cells. PAC protein expression was validated in sorted cells using whole-cell recording for PAC-specific currents.

#### *In vitro* Bacterial Clearance Assay

BMDMs were counted and cultured in 6-well plates at a density of 1.2 × 10^6^ cells per well. Infection of wildtype *E. coli* was conducted at an MOI of 20 for 1 hour at 37°C. Then the medium was changed to DMEM containing 100 μg/mL gentamycin to remove extracellular bacteria. Following 1 or 2 hours of digestion, cells were washed with ice-cold PBS and lysed with 0.3% (vol/vol) Triton X-100 for 5 minutes. Cell lysates containing undigested bacteria were serially diluted with PBS and inoculated on LB plates followed by CFU determination after overnight incubation at 37°C. The uptake of the bacteria was determined by culturing bacteria from cell lysates immediately after 1 hour of infection.

#### Measurement of Phagosome pH and Protease Activity

Analysis of phagosome pH was measured by a ratiometric method described previously (Osei-Owusu et al., 2021). Dual-labeled zymosan particles were employed for these measurements. PhrodoGreen and AF633-labeled zymosan particles were used to assess pH, while DQGreen BSA and AF633-labeled zymosan particles were employed to evaluate protease activity. The conjugation of dual-labeled zymosan particles follows the Amine-Reactive Probe Labeling Protocol (Thermo Fisher Scientific). Subsequently, BMDM or Raw264.7 cells were treated with dual-labeled zymosan particles as indicated. Single cell suspensions were obtained from detached cells using macrophage detachment buffer and subjected to FACS analysis. For phagosome pH measurement, singlet cells positive with both PhrodoGreen and AF633 were gated, and the fluorescence intensity was quantified. A standard curve was generated using the Intracellular pH Calibration Buffer kit (Thermo Fisher Scientific) with different pH values (5.5, 6.5, and 7.5) in the presence of 10 mM nigericin and valinomycin to equilibrium the pH with the calibration buffers. The phagocytic index was calculated by subtracting AF633 MFI from baseline autofluorescence value. The PhrodoGreen/AF633 ratio was used to normalize phagocytic uptake and used for eventual ratiometric calculation. For protease activity, the DQGreen BSA/AF633 ratio was used to normalize phagocytic uptake, and the value was used to compare the protease activity between experimental groups.

For pH and protease activity measurements in peritoneal cells, mice were peritoneally injected with either pH or protease dual-labeled zymosans at an MOI of 20 for 30 minutes. Peritoneal lavage was then conducted to acquire peritoneal cells, which were subjected to surface staining. FACS analysis was used to quantify MFI of PhrodoGreen/AF633 or DQGreen BSA/AF633 within gated LPM, and these values were used for further calculations.

#### Immunofluorescence staining and Confocal Microscopy

Immunofluorescence staining was performed as previously described (Osei-Owusu et al., 2021). THP-1 macrophages were differentiated on coverslips 48 hours before fixation. Isolated LPM macrophages were incubated on coverslips for 20 minutes for adherence. Peritoneal *E. coli* was collected from peritoneal fluids and stained with propidium iodide (PI, 1 μg/mL, Invitrogen) for 30 min at 37 °C before centrifugating 3000xg for 5 minutes onto coverslips followed by fixation. For phagosomal PAC, Rab5, Rab7, and Lamp1 imaging, cells were fixed with methanol for 10 minutes at −20°C followed by acetone permeabilization 1 minute at −20°C. For EEA1, Rab11, flagellin, IRF1, pSTAT1, and pp65 imaging, cells were fixed in 4% PFA for 20 minutes at room temperature (RT) followed by 0.3% Triton X-100 permeabilization 30 minutes at RT. Cells further underwent blocking by 5% bovine serum albumin (BSA) in PBS for 60 minutes at RT and incubated with indicated antibodies in 5% BSA buffer overnight at 4°C. For non-permeabilized surface F4/80 and gasdermin D staining, cells were 4% PFA fixed without permeabilization and blocked with 5% BSA for 60 minutes before primary antibody incubation. After washing with PBS three times, samples were incubated with Alexa Fluor–conjugated secondary antibodies (1:500; Invitrogen) for 1 hour and DAPI (1 μg/mL) 10 minutes at RT. Coverslips were washed with PBS prior to mounting with Aqua-poly Mount (Polysciences, Inc). Mounted samples were cured overnight at RT and imaged. Single plane or Z-stack images were taken under Zeiss LSM900 confocal microscopes with 60x objective lens. Images were analyzed with ImageJ or ZEN software.

#### Patch Clamp Electrophysiology

Whole-cell patch clamp recordings were performed as previously described (Osei-Owusu et al., 2021). Cells were recorded in an extracellular solution (ECS) containing: 145mM NaCl, 1.5mM CaCl_2_, 2mM MgCl_2_, 2mM KCl, 10mM HEPES, and 10mM glucose (pH adjusted to pH 7.3 with NaOH and osmolality was 300–310 mOsm/kg). Acidic ECS with pH 4.6 was made with the same ionic composition without HEPES but with 5mM Na_3_-citrate as buffer and the pH was adjusted using citric acid. The acidic solution was perfused using a gravity perfusion system. Recording pipettes (2–4MU) were filled with internal solution containing: 135mM CsCl, 2mM CaCl_2_, 1mM MgCl_2_, 4mM MgATP, 0.5mM Na3 -GTP, and 5mM EGTA (pH adjusted to 7.2 with CsOH and osmolality was 280–290 mOsm/kg). For hypotonicity-activated VRAC current recordings, hole-cell patch-clamp configuration was established in an isotonic bath solution containing 90 mM NaCl, 2 mM KCl, 1 mM MgCl2, 2 mM CaCl2, 10 mM Hepes, 10 mM glucose, and 100 mM mannitol (pH adjusted to pH 7.3 with NaOH and osmolality adjusted to 310 mOsm/kg), and then a hypotonic solution that has the same ionic composition but without mannitol was applied. Recording electrodes (2 to 4 megohms) were filled with a standard internal solution containing 133 mM CsCl, 10 mM Hepes, 4 mM Mg-ATP, 0.5 mM Na3–guanosine 5′ -triphosphate (GTP), 2 mM CaCl2, and 5 mM EGTA (pH was adjusted to 7.2 with CsOH, and osmolality was 290 to 300 mOsm/kg).

All recordings were done at room temperature with MultiClamp 700B amplifier and 1550B digitizer (Molecular Devices). Data acquisition was performed with pClamp 10.7 software (Molecular Device), filtered at 2 kHz and digitized at 10 kHz. Voltage ramp pulses were applied every 5 s from −100 to +100 mV at a holding potential of 0 mV. Voltage step pulses were applied every 3 s for a duration of 1 s, from −100 to +100 mV at a holding potential of 0 mV in a 20-mV increment. All data were analyzed using Clampfit 10.7 and GraphPad Prism software was used for all statistical analyses.

#### LPM RNA sequencing

Mice were subjected to peritoneal infection for 30 minutes and underwent peritoneal lavage to acquire peritoneal cells. Cells were washed with ice-cold PBS and collected in FACS buffer. Cells further underwent surface staining for LPM gating. Subsequently, Sony MA900 Multi-Application Cell Sorter was used for isolating LPM cells. 3 samples from PBS- and infection-treated mice from control or *Cx3cr1*-PAC^−/−^ mice were collected. Total RNA was extracted using the RNeasy Micro kit (Qiagen). Total RNA quality was determined by the A_260_/A_280_ and A_260_/A_230_ as well as Agilent 2100 Bioanalyzer.

RNA sequencing service was carried out by Novogene Company. In brief, libraries were constructed using the NEBNext Ultra II RNA library Prep Kit for Illumina (New England Biolabs) and sequenced on an illumina NovaSeq 6000 System. Reads were mapped to GRCm38/mm10 reference mouse genome with Hisat2 v2.0.5. FeatureCounts v1.5.0-p3 was used to count the reads numbers mapped to each gene. Fragments per kilobase per million mapped reads (FPKM) of each gene was calculated based on the length of the gene and reads count mapped to this gene. Further differential expression analysis, Gene Set Enrichment analysis, Gene Ontology analysis, and KEGG pathway analysis were carried out by clusterProfiler R package. Data were visualized using GraphPad Prism and NovoMagic (Novogene).

#### High-Dimensional Mass Cytometry (CyTOF)

Mice were subjected to peritoneal infection for 4 hours and underwent peritoneal lavage to acquire peritoneal cells. Cells were first washed with ice-cold MAXPAR PBS (Fluidigm) then stained with 1 mL of 1 μM Cell-ID cisplatin-^194^Pt (Fluidigm) for 3 minutes at RT to exclude dead cells. Next, cells were prepared and stained following Maxpar Cell Surface Staining with Fresh Fix protocol (Fluidigm). In brief, cells were Fc blocked and stained with antibody panel as provided in the Supplemental Material Table for 30 minutes at RT. A fresh fix was then conducted with incubating cells with freshly made 1.6% formaldehyde solution (Thermo Fisher Scientific) for 10 minutes at RT. Lastly, cells were transferred to intercalation solution (125 nM of Cell-ID intercalator-Ir in Maxpar Fix and Perm Buffer) before submitted to Johns Hopkins SKCCC Flow/Mass Cytometry Core for data acquisition. Of note, 168Er conjugated Tim-4 and 156Gd conjugated SIRPα antibodies were made from conjugating isotopes to Tim-4 and SIRPα antibodies (BioLegend) using Maxpar X8 antibody labeling kits (Fluidigm).

#### OMIQ analysis and workflow

cyTOF FCS files acquired and exported from the Johns Hopkins SKCCC Flow/Mass Cytometry Core were uploaded to OMIQ. Live, singlet and CD45-positive cells were gated before subsampling into a maximum equal distribution across groups. After subsampling, dimensional reduction methods (tSNE, opt-SNE, and UMAP) were performed using default parameters, followed by PhenoGraph, SPADE, and FlowSOM clustering. Data was further analyzed with EdgeR to determine significantly different clusters, or by generating heatmaps with Euclidean clustering. OMIQ was also used for performing manual gating strategies for identified populations and the quantification of population sizes. Data were visualized using OMIQ or GraphPad Prism.

#### Phagosome proteome and database transcriptome analysis

The phagosome proteome dataset was downloaded from ProteomeXchange (PXD001293). The list of proteins was curated by selecting transmembrane proteins from the reported proteome before filtering out the unknown proteins or proteins that have been reported with substrates other than ions. Additionally, proteins that were only detected in either BMDM or Raw264.7 cells or with zero signal intensity in any samples were also excluded. For the GeneCards database analysis, each of proteins within the phagosome proteome list was surveyed on the GeneCards database with corresponding gene for the subcellular localization confidence. Genes with confidence above 3 on endosomes or lysosomes were used for the following analysis. Gene expression RNA-Seq values for the Immunological Genome Project (Heng et al., 2008) (ImmGen) was downloaded from the online portal. The expression values were used for the subsequent z-score calculation across various cell types or the direct comparison between genes. Dataset for GSE134364 or GSE94916 was downloaded from the NCBI Gene Expression Omnibus (GEO). The expression values were acquired from GEO2R portal and were used for the statistical analysis.

#### Inflammasome Activation

Freshly isolated LPMs were plated at a density of 5 × 10^5^ cells/well in 12-well plates. For *E. coli* infection, bacterium at MOI 20 was simultaneously plated in 12-well plates. For ATP and nigericin activation, cells were activated by LPS (1 μg/mL) for 4 hours upon plating before ATP (4 mM) or nigericin (5 μM) treatment for 30 minutes. Subsequently, cell supernatants were collected, centrifugated at 12000xg to remove dead cells and debris, and analyzed for IL-1β, IL-18, or LDH release as indicated.

#### E64d phagolysosome cathepsin inhibition

E64d cathepsin inhibitor was acquired from Selleck Chemicals. For *in vivo* E64d inhibition in LPM cells, E64d (0.15 mg/kg) in 100 μL PBS was peritoneally injected 30 minutes before *E. coli* injection. For *in vitro* E64d inhibition, isolated LPM was pretreated with 5μM of E64d for 30 minutes before subsequent *E. coli* infection.

#### Quantitative Real-Time PCR

Total RNA was isolated using TRIzol reagent (Life Technologies). 1-2 μg of total RNA was used to generate the cDNA using the High-Capacity cDNA Reverse Transcription Kit (Applied Biosystems). The reaction was run in QuantStudio 6 Real-Time PCR Systems using 0.2 μl of cDNA in a 15 μl reaction. For the amplification of PAC mRNA, IDT PrimeTime Predesigned qPCR probes targeting mouse *Tmem206* exon 1-2 were used in combination with PerfeCTa qPCR SuperMix (Quanta Bio) according to manufacturer’s instruction. Primers targeting other genes were provided in the Supplemental Material Table and used in combination with SYBR green master mix (Applied Biosystems) according to manufacturer’s instruction. The quantification was collected as CT value. Calibrations and normalizations were performed using the 2^-ΔΔCT method. *Gapdh* or *Actb* was used as the reference gene.

#### Protein Immunoblotting

Cytosolic cell extracts were prepared using hypotonic buffer to exclude organelles and nucleus (250 mM sucrose, 20 mM HEPES, 10 mM KCl, 1.5 mM MgCl_2_, 1 mM EGTA, 1 mM EDTA, 1 mM Pefabloc). Cytosolic extracts were then precipitated by adding trichloroacetic acid (final concentration 5%). After 10 min on ice, the proteins were pelleted by centrifugation at 20,800xg for 15 min. The pellets were then resuspended in RIPA buffer (Cell Signaling Technology). Whole cell lysates of THP-1 cell or isolated LPMs were prepared in RIPA buffer with protease inhibitor cocktails (Roche). Lysates in RIPA buffer were cleared of debris by centrifugation at 12000xg for 10 min at 4°C and the concentration of protein lysates was determined by BCA assay kit (Thermo Fisher Scientific). Lysates were then collected and made into same concentrations with water and Blot 4x LDS sample buffer (Invitrogen) before boiling at 75°C for 10 min prior to SDS-PAGE electrophoresis. Novex 4-20% gradient gel (Invitrogen) was used for electrophoretic separation according to manufacturer’s instruction. For subsequent antibody detection, nitrocellulose membranes were used for protein transfer. Primary antibodies were made as indicated by manufacturers and incubated with nitrocellulose membranes overnight at 4°C. Following primary antibody incubation, samples were incubated with HRP-conjugated anti-mouse, anti-rabbit, or anti-rat secondary antibody (Proteintech) for 1 hour at RT for the visualization.

#### Statistics

All data were analyzed using GraphPad Prism software. Images were analyzed with ImageJ (U.S. National Institutes of Health) and ZEN software (ZEISS). Data were presented as mean with standard error of the mean (SEM) as indicated in figure legends. Survival curve data was presented as a Kaplan-Maier plot with a log-rank (Mantel-Cox) test used to compare susceptibility between different groups. Bar plots, dot plots, donut graphs, survival plots, volcano plots, and heatmaps were plotted using GraphPad Prism. FACS dot plots, histograms, FlowSOM trees, SPADE trees, and UMAPs were plotted using FlowJo (BD) or OMIQ. Graphical abstract and illustrations were made using Biorender.com. The sample sizes were indicated in figures and legends. Statistical significance was determined using t-tests (two-tailed), one-way ANOVA, or two-way ANOVA followed by multiple comparison tests. A *p* value less than 0.05 was considered statistically significant.

