## Supplementary Figures and legends for "Phagosome-mediated anti-bacterial immunity is governed by the proton-activated chloride channel in peritoneal macrophages"

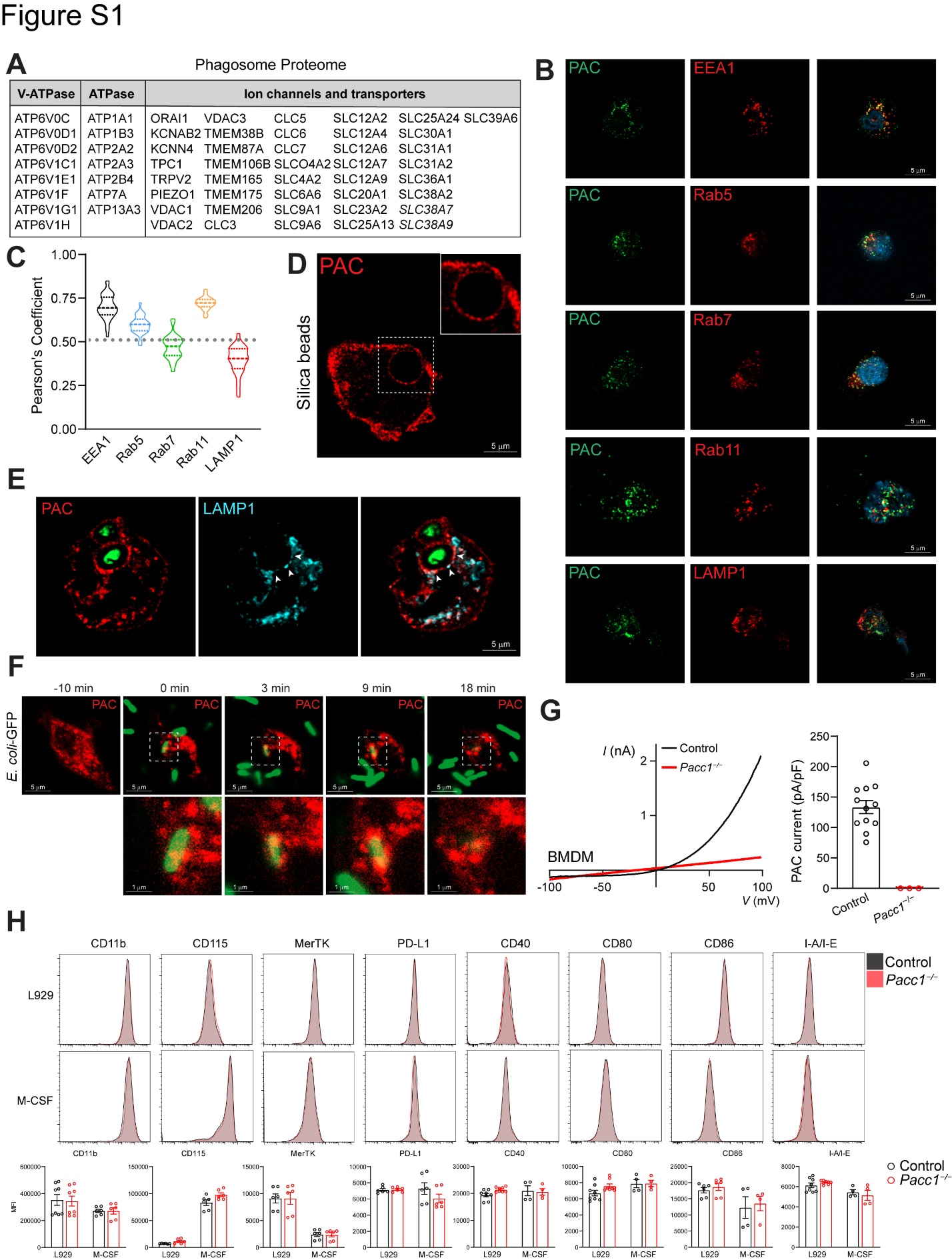
**Supplementary Figure 1. PAC is a phagosome chloride channel that is downregulated upon macrophage activation.**

**A,** List of ion channels and transporters in the phagosome proteome (ProteomeXchange: PXD001293).

**B-C**, Representative confocal images (**B**) and Pearson’s coefficient (**C**) of single-plane THP1 cells. A cutoff of 0.5 was made to indicate colocalization. Data shown as representative of at least 20 cells. Scale bar as indicated.

**D,** Representative confocal image of single-plane THP1 cells with silica beads. Data shown as representative of at least 20 cells. Scale bar as indicated.

**E,** Representative confocal image of THP1 cells with pH-sensing zymosans. The image corresponds to Fig. 1D with a single-plane to highlight LAMP1 localization. Data shown as representative of at least 20 cells. Scale bar as indicated.

**F,** Representative confocal images of frames from live-cell imaging of PAC-mCherry expressing Raw264.7 cells after *E coli-GFP* infection. Data shown as representative of at least 20 cells. Scale bar as indicated.

**G**, Representative I-V relationship (left) and current density (right) of whole-cell PAC current in BMDMs. control, n = 12 cells; KO, n = 3 cells.

**G,** FACS analysis histogram (top) and MFI quantification (bottom) of L929 conditioned medium- or M-CSF-differentiated BMDMs. Data represented from n = 8 mice.

All control groups represent *Pacc1^F/F^* genotype. Data are reported as mean ± SEM between independent experiments.

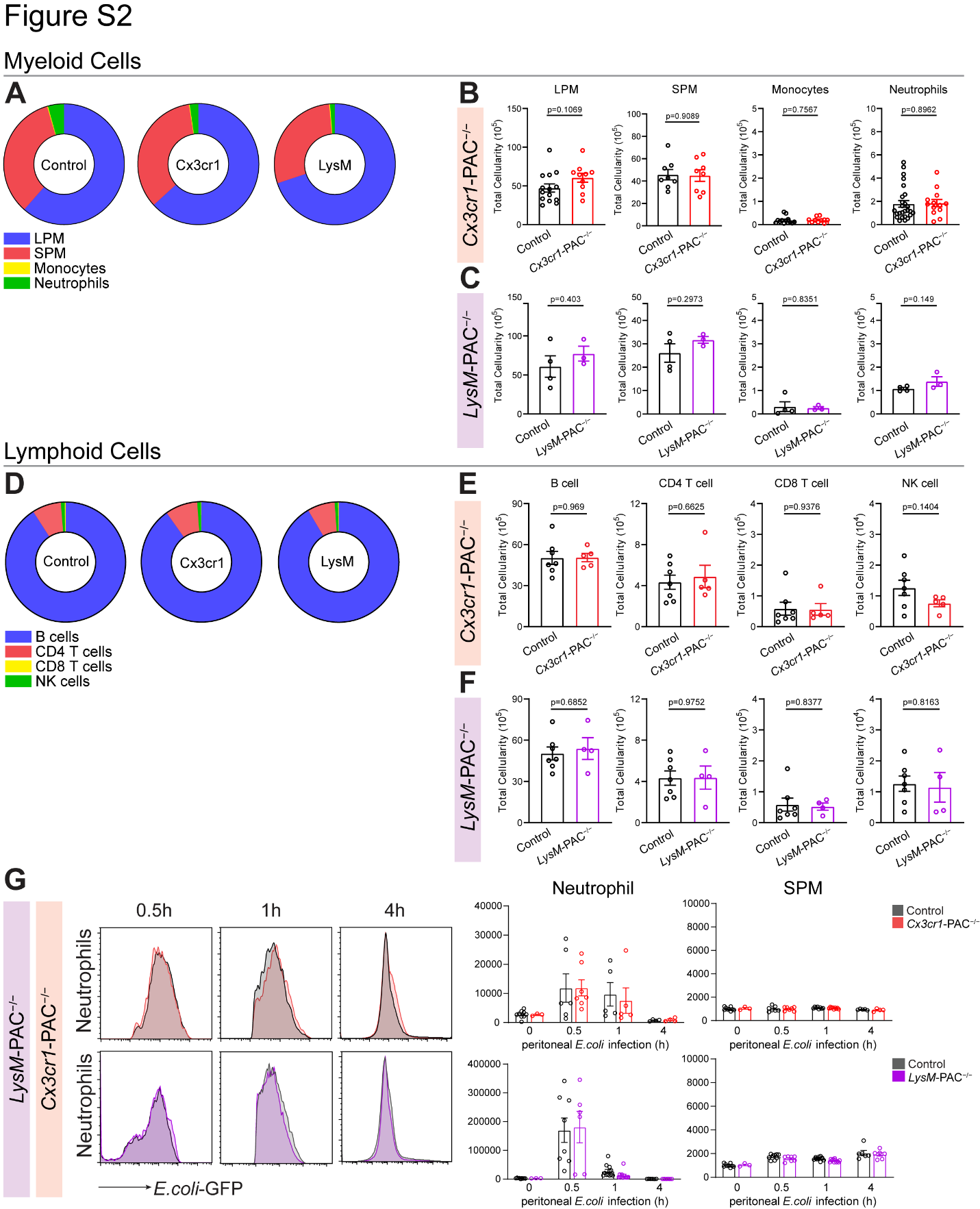
**Supplementary Figure 2. PAC deletion does alter immune cell development.**

**A**, Donut plots of peritoneal myeloid cell populations with respect to genotypes.

**B-C**, FACS quantification of myeloid cell populations in peritoneal fluids.

**D**, Donut plots of splenic lymphoid cell populations with respect to genotypes.

**E-F**, FACS quantification of splenic lymphoid cell populations.

**G**, FACS analysis histogram (left) and MFI quantification (right) of *in vivo* *E. coli*-GFP clearance assay. For histograms, *E. coli*-GFP fluorescence in LPM was plotted at indicated times. For quantifications, GFP MFI in LPM was quantified at indicated times. At least 500,000 cells were collected per condition. Data represented from n > 4 mice.

All control groups represent *Pacc1^F/F^* genotype. Data are reported as mean ± SEM between independent experiments. Unpaired t-test for **B, C, E, F**.

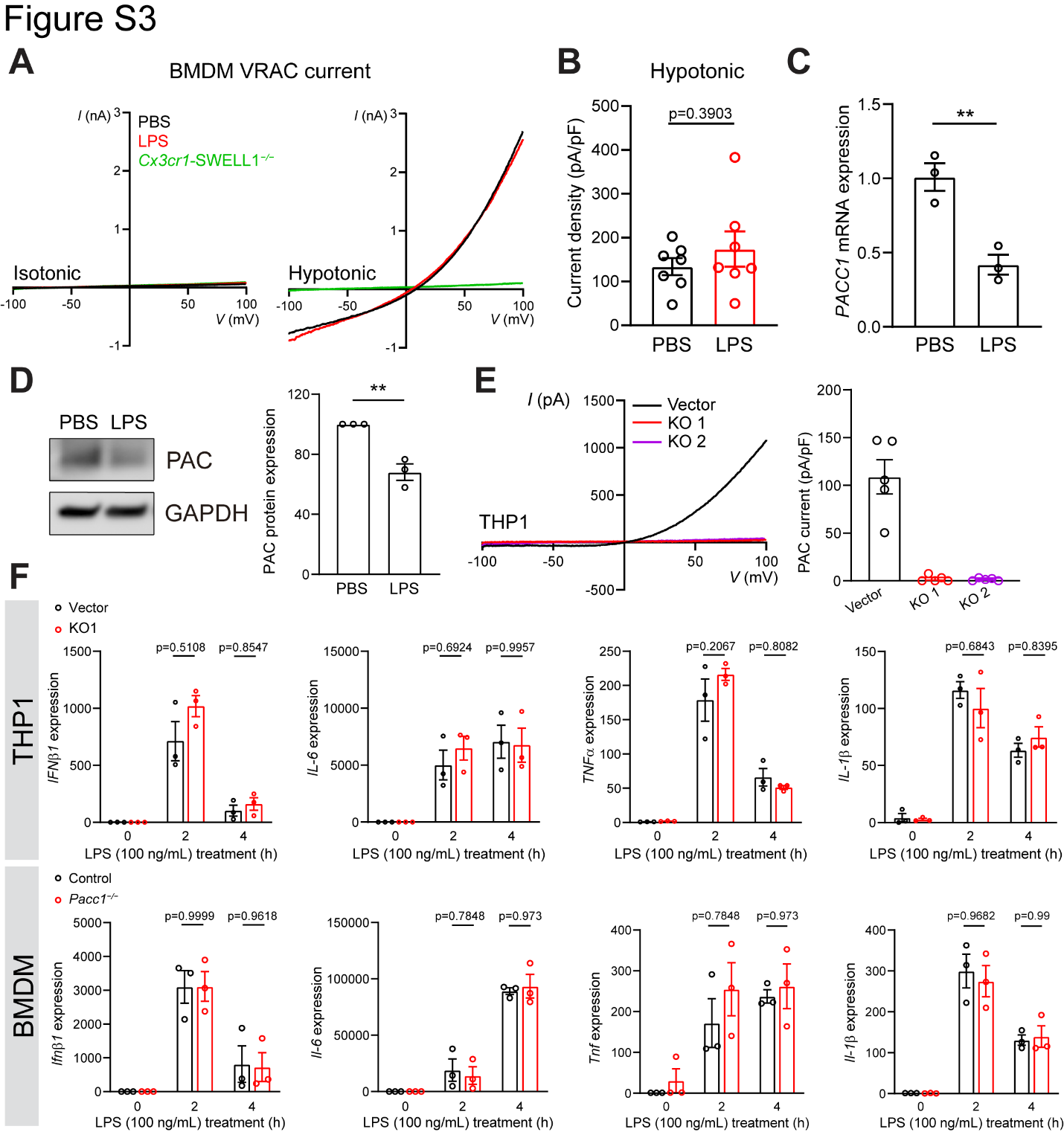
**Supplementary Figure 3. PAC does not alter LPS-induced inflammatory gene expression.**

**A-B**, Representative I-V relationship (**A**) and current density (**B**) of whole-cell VRAC current in BMDMs post 4 hours LPS (1 μg/mL) treatment. PBS: n = 7 cells from 3 mice. LPS: n = 7 cells from 3 mice.

**C**, *PACC1* mRNA expression in THP1 cells after LPS (100 ng/mL) treatment. Data represented from n = 3 independent experiments.

**D**, Immunoblotting (left) and quantification (right) of THP1 cells after LPS (100 ng/mL) treatment. Data represented from n = 3 independent experiments.

**E**, Representative I-V relationship (left) and current density (right) of whole-cell PAC current in THP1 cells. Control, n = 5 cells; KO1, n = 5 cells; KO2, n = 5 cells.

**D**, *IFNβ*, *IL-6*, *TNFα*, and *IL-1β* mRNA expression in THP1 cells (top) or *Ifnβ*, *Il-6*, *Tnfα*, and *Il-1β* mRNA expression in BMDMs (bottom) after LPS (100 ng/mL) treatments. Data represented from n = 3 mice or independent experiments.

All control groups represent *Pacc1^F/F^* genotype. Data are reported as mean ± SEM between independent experiments. Unpaired t-test for **B, C, D**. Two-way ANOVA with Sidak’s test for **F**. **p < 0.01.

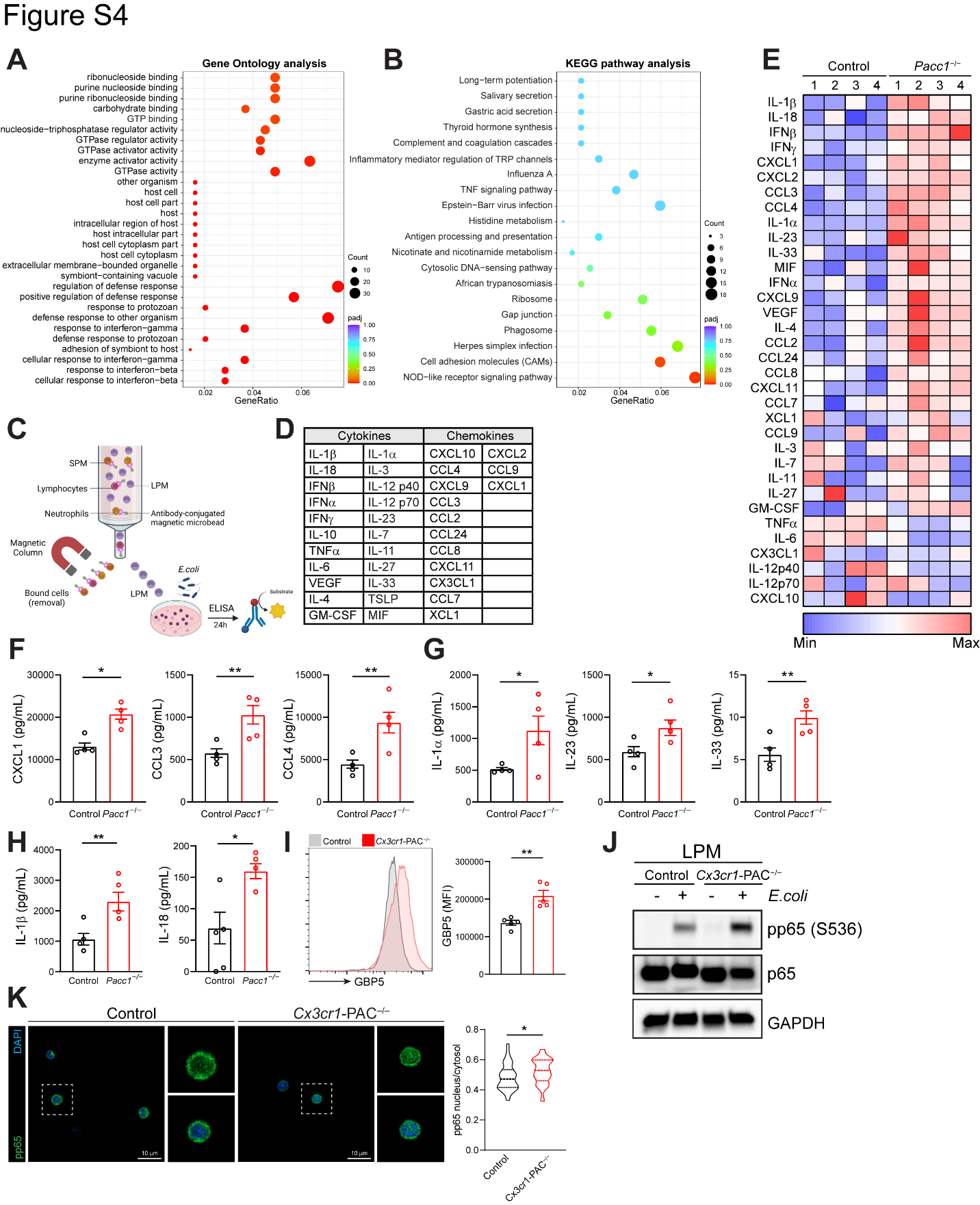
**Supplementary Figure 4. Hyperactivation of PAC-deficient LPMs after *E. coli* infection.**

**A-B**, Dot plots of analyzed GO terms (**A**) or KEGG analysis (**B**) corresponding to Fig 3B.

**C,** Schematic illustration of LPM isolation and *E. coli* infection for secretome analysis.

**D**, List of analyzed secretome.

**E**, Heatmap depicting Z-score of highlighted secreted proteins.

**F-H**, ELISA of differentially secreted proteins in (**E**), including chemokines (**F**), alarmins (**G**), and IL-1β and IL-18 (**H**). Data represented from n = 4 mice.

**I**, FACS analysis histogram (left) and MFI quantification (right) of LPMs after 30 min of peritoneal infection. Data represented from n = 5 mice.

**J**, Immunoblotting of MACS-isolated LPM. LPMs from 2 mice 15 min post peritoneal infection were pooled and used in 1 experiment. Data represented from n = 3 independent experiments.

**K**, Representative confocal images (left) in single-plane and quantification (right) of MACS-isolated LPMs 30 min post peritoneal infection. Control LPM, n = 52 cells; *Cx3cr1*-PAC^−/−^ LPM, n = 42 cells.

All control groups represent *Pacc1^F/F^* genotype. Data are reported as mean ± SEM between independent experiments. Unpaired t-test for **F, G, H, I** and **K**. *p < 0.05, **p < 0.01.

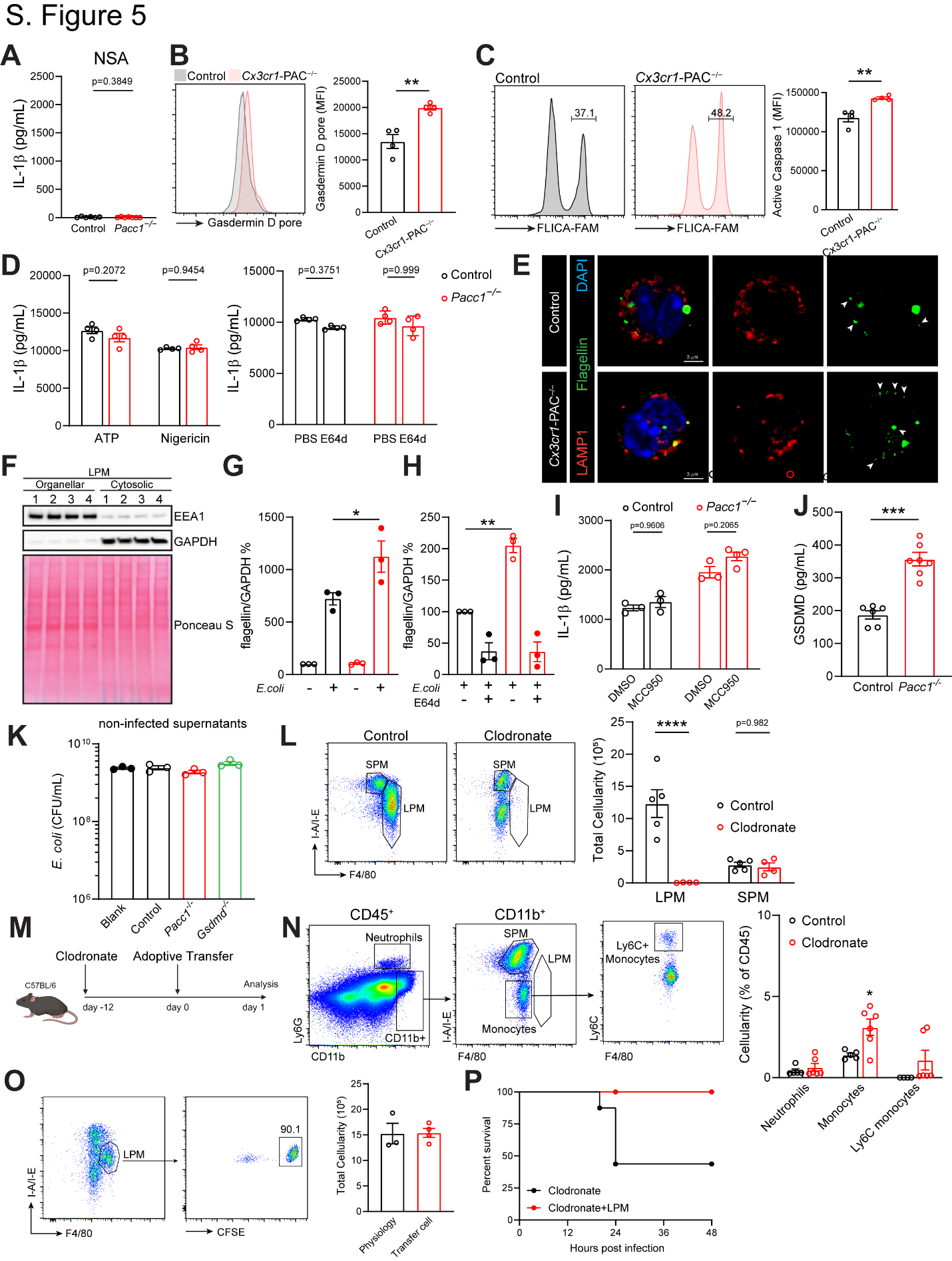
**Supplementary Figure 5. Augmented phagosome bacterial degradation facilitates inflammasome activation and pyroptosis in PAC-deficient LPMs.**

**A**, ELISA of IL-1β from MACS-isolated LPMs pretreated with NSA (10μM) for 30 min following *E. coli* infection *in vitro*. Data represented from n = 5 mice.

**B**, FACS analysis histogram (left) and MFI quantification (right) of LPMs after 30 min of peritoneal infection. Data represented from n = 4 mice.

**C**, FACS analysis histogram (left) and MFI quantification (right) of LPMs after 30 min of peritoneal infection. Data represented from n = 4 mice.

**D**, ELISA of IL-1β from MACS-isolated LPMs primed with LPS (1 μg/mL) for 4 hours followed by ATP (4 mM) or nigericin (10 μM) for 30 minutes (left) or E64d pretreatment before LPS priming and ATP challenge (right). Data represented from n = 4 mice.

**E**, Representative confocal images of MACS-isolated LPMs after 60 min of peritoneal infection. Data represented from n > 20 cells.

**F**, Immunoblotting of MACS-isolated LPM organellar and cytosolic extraction. LPMs from 2 mice were pooled and used in 1 experiment. Data represented from n = 4 independent experiments.

**G-H**, Immunoblotting quantification for Fig. 4H (**G**) and Fig. 4I (**H**).

**I**, ELISA of IL-1β from MACS-isolated LPMs pretreated with MCC950 (100nM) for 30 min following *E. coli* infection *in vitro*. Data represented from n = 3 mice.

**J**, ELISA of gasdermin D from MACS-isolated LPMs after *E. coli* infection *in vitro*. Data represented from n > 6 mice.

**K**, *E. coli* cultured with antibiotic-free supernatants, collected from non-infected control, *Pacc1^−/−^*, or *Gsdmd^−/−^* LPMs, for 6 h at 37C. Blank, antibiotic-free RPMI medium.

**L**, FACS analysis (left) and quantification (right) of peritoneal cells 12 days after clodronate-liposome injection. Data represented from n > 4 mice.

**M**, Schematic illustration of depletion-adoptive transfer model without peritoneal infection

**N**, FACS analysis (left) and quantification (right) of peritoneal cells 12 days after clodronate-liposome injection. Data represented from n > 4 mice.

**O**, FACS analysis (left) and quantification (right) of CellTrace^+^ LPM 24 hours after adoptive transfer. Data represented from n = 3 mice.

**P**, Survival of mice after *E. coli* peritoneal infection at 10^7^ CFU/mL; Clodronate n = 8. Clodronate+LPM n = 9.

Control represents *Pacc1^F/F^* genotype in **A-K** or control liposomes in **L-N**. Data are reported as mean ± SEM between independent experiments. Unpaired t-test for **A, B, C, J, L.** Two-way ANOVA with Sidak’s test for **D, G, H, I**. *p < 0.05, **p < 0.01, ***p < 0.001, ****p < 0.0001.
