## Supplementary material table for "Phagosome-mediated anti-bacterial immunity is governed by the proton-activated chloride channel in peritoneal macrophages"

| **REAGENT or RESOURCE** | **SOURCE** | **IDENTIFIER** |
| --- | --- | --- |
| **Antibodies** | | |
| Mouse anti-PAC | This study | N/A |
| Rabbit anti-EEA1 | Cell Signaling Technology | Cat # 3288; RRID: AB_2096811 |
| Rabbit anti-Rab5 | Cell Signaling Technology | Cat # 3547; RRID:AB_2300649 |
| Rabbit anti-Rab7 | Cell Signaling Technology | Cat # 9367; RRID:AB_1904103 |
| Rabbit anti-Rab11 | Cell Signaling Technology | Cat # 5589; RRID:AB_10693925 |
| Rabbit anti-LAMP1 | Cell Signaling Technology | Cat # 9091: RRID: AB_2687579 |
| Rabbit anti-GAPDH | Cell Signaling Technology | Cat # 2118; RRID: AB_561053 |
| Alexa Fluor 488 goat anti-rabbit IgG | Thermo Fisher Scientific | Cat # A-11008; RRID: AB_143165 |
| Alexa Fluor 488 goat anti-mouse IgG | Thermo Fisher Scientific | Cat # A28175; RRID:AB_2536161 |
| Alexa Fluor 546 goat anti-mouse IgG | Thermo Fisher Scientific | Cat # A-11030; RRID:AB_2534089 |
| Alexa Fluor 647 goat anti-mouse IgG | Thermo Fisher Scientific | Cat # A-21235; RRID:AB_2535804 |
| Rabbit anti-goat IgG HRP-conjugated | R&D systems | Cat # HAF017; RRID: AB_562588 |
| HRP-conjugated Affinipure Goat Anti-Mouse IgG(H+L) | Proteintech | Cat # SA00001-1; RRID:AB_2722565 |
| HRP-conjugated Affinipure Goat Anti-Rabbit IgG(H+L) | Proteintech | Cat # SA00001-2; RRID:AB_2722564 |
| Rabbit anti-NLRP3 | Cell Signaling Technology | Cat # 15101; RRID:AB_2722591 |
| Rabbit anti-cleaved Gasdermin D | Cell Signaling Technology | Cat # 10137; RRID:AB_2923068 |
| Gasdermin-D, rabbit monoclonal, clone EPR20859 | Abcam | Cat # ab219800; RRID:AB_2888940 |
| GSDMD N-Terminal Antibody(Mouse specific) | Affinity Biosciences | Cat # DF13758; RRID: AB_3076218 |
| Mouse monoclonal anti-caspase-1 | Adipogen | Cat # AG-20B-0042; RRID: AB_2490248 |
| Rabbit polyclonal anti-GBP2 | Proteintech | Cat # 11854-1-AP; RRID: AB_2109336 |
| Rabbit polyclonal anti-GBP5 | Proteintech | Cat # 13220-1-AP; RRID: AB_2109348 |
| Rabbit monoclonal anti-IRF1 | Cell Signaling Technology | Cat # 8478S; RRID: AB_10949108 |
| Rabbit polyclonal anti-Flagellin | Abcam | Cat #ab93713; RRID:AB_10563522 |
| Rabbit monoclonal anti- Phospho-STING (Ser365) | Cell Signaling Technology | Cat # 72971S; RRID: AB_2799831 |
| Rabbit monoclonal anti-STING | Cell Signaling Technology | Cat # 13647S; RRID: AB_ 2732796 |
| Rabbit monoclonal anti- Phospho-IRF3 (Ser396) | Cell Signaling Technology | Cat # 29047S; RRID: AB_2773013 |
| Rabbit monoclonal anti-IRF3 | Cell Signaling Technology | Cat # 4302S; RRID: AB_ 1904036 |
| Rabbit monoclonal anti- Phospho-STAT1 (Tyr701) | Cell Signaling Technology | Cat # 9167S; RRID: AB_ 561284 |
| Rabbit monoclonal anti-STAT1 | Cell Signaling Technology | Cat # 9172S; RRID: AB_ 2198300 |
| Rabbit polyclonal anti-OAS1/3 | Proteintech | Cat # 14955-1-AP; RRID: AB_2158292 |
| Rabbit polyclonal anti-OAS2 | Proteintech | Cat # 19279-1-AP; RRID: AB_10642832 |
| Rabbit monoclonal anti- Phospho-p65 (Ser536) | Invitrogen | Cat # MA5-15160; RRID: AB_ 10983078 |
| Rabbit monoclonal anti-p65 | Cell Signaling Technology | Cat # 8242S; RRID: AB_ 10859369 |
| Antibodies for Flow Cytometry | | |
| TrueStain FcX anti-CD16/32 antibody | BioLegend | Cat # 156603; RRID:AB_2783137 |
| Anti-mouse/human CD11b FITC clone M1/70 | BioLegend | Cat # 101206; RRID:AB_312789 |
| Anti-mouse CD115 PE/Dazzle594 clone AFS98 | BioLegend | Cat # 135527; RRID:AB_2566522 |
| Anti-mouse CD40 PE/Cy7 clone 3/23 | BioLegend | Cat # 124621; RRID:AB_10933422 |
| Anti-mouse CD80 APC clone 16-10A1 | eBioscience | Cat # 17-0801-81;  RRID:AB_469416 |
| Anti-mouse MerTK APC/Cy7 clone 2B10C42 | BioLegend | Cat # 151519; RRID:AB_2876507 |
| Anti-mouse CD274 (B7-H1, PD-L1) BV421 clone 10F.9G2 | BioLegend | Cat # 124315; RRID:AB_10897097 |
| Anti-mouse CD86 BV605 clone GL-1 | BioLegend | Cat # 105037; RRID:AB_11204429 |
| Anti-mouse I-A/I-E BV785 clone M5/114.15.2 | BioLegend | Cat # 107645; RRID:AB_2565977 |
| Anti-mouse TLR4 APC clone SA15-21 | BioLegend | Cat # 145405; RRID:AB_2562502 |
| Anti-mouse CD14 BV421 clone SA14-2 | BioLegend | Cat # 123329; RRID:AB_2721526 |
| Anti-mouse CD45 APC/Cy7 clone 30-F11 | BioLegend | Cat # 103116; RRID:AB_312981 |
| Anti-mouse/human CD11b BV421 clone M1/70 | BioLegend | Cat # 101251; RRID:AB_2562904 |
| Anti-mouse Ly6G BV605 clone 1A8 | BioLegend | Cat # 127639; RRID:AB_2565880 |
| Anti-mouse F4/80 PE clone BM8 | BioLegend | Cat # 123110; AB_893486 |
| Anti-mouse/human CD11b PE/Dazzle594 clone M1/70 | BioLegend | Cat # 101256; RRID:AB_2563648 |
| Anti-mouse Ly6C APC clone HK1.4 | BioLegend | Cat # 128016; RRID:AB_1732076 |
| Anti-mouse Ly6G BV421 clone 1A8 | BioLegend | Cat # 127628; RRID:AB_2562567 |
| Anti-mouse F4/80 BV605 clone BM8 | BioLegend | Cat # 123133; RRID:AB_2562305 |
| Anti-Mouse CD4 AF700 clone GK1.5 | BioLegend | Cat # 100429; RRID:AB_493698 |
| Anti-Mouse CD8a APC clone 53-6.7 | BioLegend | Cat # 100712; RRID:AB_312751 |
| Anti-Mouse CD3e BV605 clone 145-2C11 | BioLegend | Cat # 100351; RRID:AB_2565842 |
| Anti-Mouse CD19 FITC clone 6D5 | BioLegend | Cat # 115506; RRID:AB_313641 |
| Anti-MouseCD49b PerCP-Cy5.5 clone HMα2 | BioLegend | Cat # 103520; RRID:AB_2566105 |
| Anti-Mouse NK1.1 BV421 clone PK136 | BioLegend | Cat # 108731; RRID:AB_10895916 |
| Anti-Mouse Ly6G BV785 clone 1A8 | BioLegend | Cat # 127645; RRID:AB_2566317 |
| Anti-Mouse F4/80 BV785 clone BM8 | BioLegend | Cat # 123141; RRID:AB_2563667 |
| Anti-Mouse SiglecF BV786 E50-2440 | BD Bioscience | Cat # 740956; RRID:AB_2740581 |
| Anti-Mouse/human CD11b BV785 clone M1/70 | BioLegend | Cat # 101243; RRID:AB_2561373 |
| Anti-mouse/human CD11b PE clone M1/70 | BioLegend | Cat # 101207; RRID:AB_312790 |
| Antibodies for Mass Cytometry | | |
| Anti-mouse CD45 89Y clone 30-F11 | Fluidigm / Standard BioTools | Cat # 3089005B; RRID:AB_2651152 |
| Anti-mouse CD8a 111Cd clone 53-6.7 | Fluidigm / Standard BioTools | Cat # 92J009111 |
| Anti-mouse CD4 116Cd clone RM4-5 | Fluidigm / Standard BioTools | Cat # 92J004116 |
| Anti-mouse Ly6G 141Pr clone 1A8 | Fluidigm / Standard BioTools | Cat # 3141008B; RRID:AB_2814678 |
| Anti-mouse CD11c 142Nd clone N418 | Fluidigm / Standard BioTools | Cat # 3142003B; RRID:AB_2814737 |
| Anti-mouse TCRb 143Nd clone H57-597 | Fluidigm / Standard BioTools | Cat # 3143010B |
| Anti-mouse CD3e 148Nd clone 145-2C11 | Fluidigm / Standard BioTools | Cat # 92J003148 |
| Anti-mouse CD19 149Sm clone 6D5 | Fluidigm / Standard BioTools | Cat # 3149002B; RRID:AB_2814679 |
| Anti-mouse IgM 151Eu clone RMM-1 | Fluidigm / Standard BioTools | Cat # 3151006B |
| Anti-mouse CD49b 152Sm clone DX5 | Fluidigm / Standard BioTools | Cat # 92J008152 |
| Anti-mouse CX3CR1 155Gd clone SA011F11 | Fluidigm / Standard BioTools | Cat # 92J020155 |
| Anti-mouse CD172a(SIRPa) clone P84 | BioLegend | Cat # 144001; RRID:AB_11203723 |
| Anti-mouse CD80/B7-1 158Gd clone 16-10A1 | Fluidigm / Standard BioTools | Cat # 92J023158 |
| Anti-mouse TCRgd 159Tb clone GL3 | Fluidigm / Standard BioTools | Cat # 3159012B; RRID:AB_2922919 |
| Anti-mouse CD45R(B220) 160Gd clone RA3-6B2 | Fluidigm / Standard BioTools | Cat # 3160012B |
| Anti-mouse CD117(c-kit) 166Er clone 2B8 | Fluidigm / Standard BioTools | Cat # 3166004B; RRID:AB_2801435 |
| Anti-mouse Tim-4 clone RMT4-54 | BioLegend | Cat # 130002; RRID:AB_1227802 |
| Anti-mouse CD206(MMR) 169Tm clone C068C2 | Fluidigm / Standard BioTools | Cat # 3169021B; RRID:AB_2832249 |
| Anti-mouse CD161(NK1.1) 170Er clone PK136 | Fluidigm / Standard BioTools | Cat # 3170002B; RRID:AB_2885023 |
| Anti-mouse CD86 172Yb clone A17199A | Fluidigm / Standard BioTools | Cat # 92J022172 |
| Anti-mouse F4/80 173Yb clone BM8 | Fluidigm / Standard BioTools | Cat # 92J019173 |
| Anti-mouse XCR1 175Lu clone ZET | Fluidigm / Standard BioTools | Cat # 92J018175 |
| Anti-mouse FceR1a 176Yb clone MAR-1 | Fluidigm / Standard BioTools | Cat # 3176006B; RRID:AB_2922925 |
| Anti-mouse CD11b 196Pt clone M1/70 | Fluidigm / Standard BioTools | Cat # 92J001196 |
| Anti-mouse Ly6C 198Pt clone HK1.4 | Fluidigm / Standard BioTools | Cat # 92J010198 |
| Anti-mouse I-A/I-E 209Bi clone M5/114.15.2 | Fluidigm / Standard BioTools | Cat # 3209006B; RRID:AB_2885025 |
| Bacterial and virus strains | | |
| *Escherichia coli* (Migula) Castellani and Chalmers | ATCC | Cat # 25922 |
| *Escherichia coli* GFP | ATCC | Cat # 25922GFP |
| *Escherichia coli* (Migula) Castellani and Chalmers  (Resistant to ertapenem and imipenem) | ATCC | Cat # BAA-2469 |
| **Chemicals, peptides, and recombinant proteins** | | |
| Intracellular pH Calibration Buffer Kit | Life Technologies | Cat # 1787810 |
| SILICA BEAD IN AQUEOUS SUSPENSION | Bangs Laboratories | Cat # SS05003 |
| Clodronate Liposomes & Control Liposomes | Liposoma | Cat # CP-005-005 |
| E64d | Selleck Chemicals | Cat # S7393 |
| MCC950 | MedChemExpress | Cat # HY-12815 |
| TAK-242 | MedChemExpress | Cat # HY-11109 |
| Propidium Iodide | Thermo Fisher Scientific | Cat # P1304MP |
| Paraformaldehyde | Thermo Fisher Scientific | Cat # 416785000 |
| Adenosine 5′-triphosphate magnesium salt | Sigma | Cat # A9187 |
| Nigericin | Sigma | Cat # N7143 |
| Triton X-100 | Fisher Scientific | Cat # BP151-100 |
| RIPA buffer | Cell Signaling Technology | Cat # 9806S |
| cOmplete™ Protease Inhibitor Cocktail | Roche | Cat # 11697498001 |
| Zymosan bioparticles, unlabeled | Invitrogen | Cat # Z2849 |
| pHrodo™ Green AM Intracellular pH Indicator Dyes | Invitrogen | Cat # P35373 |
| DQ™ Green BSA | Invitrogen | Cat # D12050 |
| Alexa Fluor™ 633 NHS Ester | Invitrogen | Cat # A20005 |
| Recombinant murine M-CSF | Peprotech | Cat # 315-02 |
| Lipopolysaccharide (LPS) | Sigma | Cat # L2630 |
| DAPI | Sigma | Cat # 28718-90-3 |
| Tryptic Soy Broth | Millipore | Cat # 22092 |
| DMEM High Glu w/Gl w/ Pyr | Gibco | Cat # 11995-065 |
| RPMI 1640 w/ Glutamax | Gibco | Cat # 61870036 |
| 100 X Penicilin-Streptomycin | Gibco | Cat # 15140-122 |
| MEM Non-Essential Amino Acids Solution (100X) | Gibco | Cat # 11140050 |
| TRIzol reagent | Thermo Fisher Scientific | Cat # 15596018 |
| Blot 4x LDS sample buffer | Invitrogen | Cat # B0007 |
| LIVE/DEAD Fixable Dead Cell Aqua stain | Invitrogen | Cat # L34957 |
| Carbamacilin | Sigma | Cat # 205805 |
| Imipenem | Sigma | Cat # I0160 |
| Gentamycin | Sigma | Cat # G1264 |
| SuperSignal™ West Atto Ultimate Sensitivity Substrate | Thermo Fisher Scientific | Cat # A38555 |
| Immobilon Western Chemiluminescent HRP Substrate | Millipore | Cat # WBKLS0500 |
| Cell-ID cisplatin-^194^Pt | Fluidigm / Standard BioTools | Cat # 201194 |
| Cell-ID intercalator-Ir | Fluidigm / Standard BioTools | Cat # 201192A |
| **Critical commercial assays** | | |
| Pierce™ BCA Protein Assay Kits | Thermo Fisher Scientific | Cat # 23225 |
| High-capacity cDNA Reverse Transcription kit | Applied Biosystems | Cat # 4368814 |
| Novex 4-20% gradient gel | Thermo Fisher Scientific | Cat # XP04200BOX |
| FITC Annexin V Apoptosis Detection Kit with 7-AAD | BioLegend | Cat # 640922 |
| RNeasy Plus Mini kit | Qiagen | Cat # 74034 |
| Cyto-Fast™ Fix/Perm Buffer Set | BioLegend | Cat # 426803 |
| FAM FLICA™ Caspase-1 Kit | Bio-Rad | Cat # ICT097 |
| LegendPlex Mouse Cytokine Release Syndrome Panel (13-plex) | BioLegend | Cat # 741024 |
| LegendPlex Mouse Cytokine Panel 2 (13-plex) | BioLegend | Cat # 740134 |
| LegendPlex Mouse Proinflammatory Chemokine Panel 2 (8-plex) | BioLegend | Cat # 741067 |
| ELISA MAX™ Standard Set Mouse IL-1β | BioLegend | Cat # 432601 |
| LEGEND MAX™ Mouse MIF ELISA Kit | BioLegend | Cat # 444107 |
| Mouse IL-18 DuoSet ELISA | R&D Systems | Cat # DY7625-05 |
| Mouse CXCL1/KC DuoSet ELISA | R&D Systems | Cat # DY453-05 |
| Mouse GSDMD ELISA Kit | Abcam | Cat # ab233627 |
| Mouse IFN-beta DuoSet ELISA | R&D Systems | Cat # DY8234-05 |
| CyQUANT™ LDH Cytotoxicity Assay | Invitrogen | Cat # C20300 |
| CellTrace™ Violet Cell Proliferation Kit | Invitrogen | Cat # C34557 |
| CellTrace™ CFSE Cell Proliferation Kit | Invitrogen | Cat # C34554 |
| MACS Macrophage Isolation Kit (Peritoneum), mouse | Miltenyi Biotec | Cat # 130-110-434 |
| Maxpar® X8 Antibody Labeling Kit,156Gd—4 Rxn | Fluidigm / Standard BioTools | Cat # 201156A |
| Maxpar® X8 Antibody Labeling Kit,168Er—4 Rxn | Fluidigm / Standard BioTools | Cat # 201168A |
| Maxpar® Fix and Perm Buffer | Fluidigm / Standard BioTools | Cat # 201067 |
| Deposited data | | |
| RNA-seq data | NCBI GEO | GSE254043 |
| **Experimental models: Cell lines** | | |
| THP-1 | Dr. Andrea Cox (Johns Hopkins U) | N/A |
| PAC KO THP-1 gRNA1 | This study | N/A |
| PAC KO THP-1 gRNA1 | This study | N/A |
| Raw264.7 | ATCC | Cat # TIB-71; RRID:CVCL_0493 |
| mCherry-tagged PAC OE Raw264.7 | This study | N/A |
| PAC OE Raw264.7 | This study | N/A |
| PAC YXXL OE Raw264.7 | This study | N/A |
| L929 | ATCC | Cat # CCL-1; RRID:CVCL_0462 |
| **Experimental models: Organisms/strains** | | |
| CX3CR1-Cre [B6J.B6N(Cg)-Cx3cr1^tm1.1(cre)Jung^/J] | The Jackson Laboratory | Cat # 025524; RRID:IMSR_JAX:025524 |
| LysM-Cre [B6.129P2-Lyz2^tm1(cre)Ifo^/J] | The Jackson Laboratory | Cat # 004781; RRID:IMSR_JAX:004781 |
| Gsdmd^−/−^ (C57BL/6N-Gsdmdem4Fcw/J) | The Jackson Laboratory | Cat # 032410; RRID: IMSR_JAX:032410 |
| Casp1^−/−^ | Dr. Nathan Archer  (Johns Hopkins U) | N/A |
| PAC*^F/F^* | This study | N/A |
| CX3CR1-Cre; PAC*^F/F^* | This study | N/A |
| LysM-Cre; PAC*^F/F^* | This study | N/A |
| **Oligonucleotides** | | |
| RT-PCR primer for mIFNb1:  F: 5’-AGCTCCAAGAAAGGACGAACAT-3’  R: 5’-GCCCTGTAGGTGAGGTTGATCT-3’ | PrimerBank | N/A |
| RT-PCR primer for mIL-6:  F: 5’-CTGCAAGAGACTTCCATCCAG-3’  R: 5’-AGTGGTATAGACAGGTCTGTTGG-3’ | PrimerBank | N/A |
| RT-PCR primer for mTNFα:  F: 5’-CCCCAAAGGGATGAGAAGTT-3’  R: 5’-CACTTGGTGGTTTGCTACGA-3’ | PrimerBank | N/A |
| RT-PCR primer for mIL-1β:  F: 5’-GCAACTGTTCCTGAACTCAACT-3’  R: 5’-ATCTTTTGGGGTCCGTCAACT-3’ | *Cell Death Dis* 9, 24 (2018). | N/A |
| RT-PCR primer for hIFNb1:  F: 5’-GCTTGGATTCCTACAAAGAAGCA-3’  R: 5’-ATAGATGGTCAATGCGGCGTC-3’ | PrimerBank | N/A |
| RT-PCR primer for hIL-6:  F: 5’-ACTCACCTCTTCAGAACGAATTG-3’  R: 5’-CCATCTTTGGAAGGTTCAGGTTG-3’ | PrimerBank | N/A |
| RT-PCR primer for hTNFα:  F: 5’-CCTCTCTCTAATCAGCCCTCTG-3’  R: 5’-GAGGACCTGGGAGTAGATGAG-3’ | PrimerBank | N/A |
| RT-PCR primer for hIL-1β:  F: 5’-ATGATGGCTTATTACAGTGGCAA-3’  R: 5’-GTCGGAGATTCGTAGCTGGA-3’ | PrimerBank | N/A |
| **Software and algorithms** | | |
| GraphPad Prism 8 | GraphPad | https://www.graphpad.com/scientific-software/prism/ |
| FlowJo | BD | https://www.flowjo.com/ |
| OMIQ | OMIQ | https://www.omiq.ai/ |
| ImageJ | NIH | https://fiji.sc/ |
| Illustrator | Adobe | https://www.adobe.com/products |
| ZEN | Zeiss | https://www.zeiss.com/microscopy/en/products/software/zeiss-zen-lite.html |
| **Other** | | |
| CytoFLEX LX | Beckman Coulter | N/A |
| LSM900 | Zeiss | N/A |
| MA900 | Sony | N/A |
| Helios | Fluidigm / Standard Biotools | N/A |
| Infinite M Plex | Tecan | N/A |
